# A *Tenebrio molitor* model for *Talaromyces marneffei* infection and the importance of host cues for dimorphic switching

**DOI:** 10.64898/2026.07.29.741241

**Authors:** Bridget E. Walker, Patricia R. Jusuf, Alex Andrianopoulos

## Abstract

There is a growing body of research demonstrating the utility of invertebrate models for studies of fungal pathogenesis. In this study we evaluated *Tenebrio molitor* larvae as a model for studying talaromycosis, the infection caused by *Talaromyces marneffei. T. marneffei* is a thermally dimorphic opportunistic pathogen of humans, which transitions from hyphae to yeast when exposed to human body temperature (37°C). Using a combination of virulence assays and histopathology techniques, we have found *T. molitor* larvae to be useful simple hosts for modelling the yeast-associated disease at 37°C, as well as for studying the influence of temperature on *T. marneffei* biology. Infection establishment was found to be temperature-dependent: Larval infections could be established at both 37°C and 25°C, however 10-fold higher doses of conidia were required to cause significant disease at 25°C. Infections were also established more quickly when directly injecting larvae with the yeast cells, indicating that the yeast form has an increased capacity for host damage. *T. marneffei* in *vivo* yeast cell development was observed within larval tissues and hemolymph primarily at 37°C, but also at 25°C along with filamentous hyphal growth. The larval host environment therefore strongly supports yeast development, even partially in the absence of the 37°C signal, emphasising the combined importance of temperature and host environment for maintenance of the pathogenic morphology. This work provides a new model to assist future studies of this neglected tropical disease and improves our understanding of the complex relationship between morphology and pathogenicity in *T. marneffei*.

## Introduction

Research into human fungal infections, and ultimately the future development of novel therapeutic agents to treat these infections, relies on the availability of suitable infection models. Depending on the question being addressed, different infection models will have benefits over others for understanding of the pathogen, host and their interaction. Mammals, such as mice (*Mus musculus*) and rabbits (*Oryctolagus cuniculus*), are widely used as the model hosts for these human disease studies, although a variety of non-mammalian animal hosts also provide valuable insight into human pathogens. For instance, zebrafish (*Danio rerio*) and Arabian killifish (*Aphanius dispar*) have been shown to be useful vertebrate infection models, particularly as they are transparent as embryos, which allows for live imaging of the infection process [1, 2]. Invertebrates are now also becoming increasingly popular alternative hosts. This is largely due to ethical concerns regarding the use of vertebrates in scientific research but also advantageous for considerations of cost, sample size and handling regulations.

A range of invertebrate hosts have been used for studying host-fungi interactions [3–5]. These hosts are used mostly for studying their natural pathogens, entomopathogenic and nematophagous fungi, although several of these hosts are also susceptible to, and have been used for, studying human fungal pathogens [6]. Several components of the innate immune response are conserved between invertebrates and mammals, such as the Toll and immune deficiency (IMD) signalling pathways that trigger the production of antimicrobial peptides, as well as circulating immune cells capable of phagocytosing or encapsulating invading pathogens [7–9]. This conservation of innate immunity makes invertebrates suitable simple hosts for studying these non-native pathogens, particularly opportunistic pathogens which can cause significant disease in hosts deficient in adaptive immune responses. The relevance of these models to mammalian infection have been demonstrated for a number of pathogenic fungi, including *Aspergillus fumigatus* and *Candida albicans*, two of the most prevalent human fungal pathogens, where several read-outs were consistent between moth larvae (*Galleria mellonella*) and mice [10, 11]. These invertebrate models have additional advantages such as short lifespans and ease of maintenance, which allows researchers to screen fungal virulence quickly and in large sample groups.

*Talaromyces marneffei* is a thermally dimorphic fungal pathogen that has emerged as a significant threat to human health [12]. The fungus causes fatal systemic mycoses in immunocompromised individuals and is endemic to Thailand, Vietnam, Myanmar, Southern China, Hong Kong, Taiwan and North-eastern India [13, 14]. Despite thermal dimorphism being a common feature of fungal pathogens, *T. marneffei* is both the only dimorphic species and the only human pathogen of its genus, suggesting that its yeast morphology and ability to cause disease in humans is a result of convergent evolution [15]. Given that these traits have arisen independently, what is known about dimorphic switching in other fungal pathogens may not necessarily be applicable to *T. marneffei*, making studies of this unique pathogen’s growth and virulence critical. The developmental trajectory of a *T. marneffei* conidium is dependent on both temperature and host factors [16]. At 25°C, *T. marneffei* conidia germinate to produce multicellular hyphae, while inside mammalian and fish host cells (at 37°C and 33°C respectively), conidia germinate into unicellular yeast cells which then continue to proliferate [17, 18]. At 37°C *in vitro T. marneffei* conidia do not germinate directly into yeast cells. Instead, they initially produce a highly-branched hyphal form that produces uninucleate cells after ~48 hours, which ‘break-off’ from the hyphal form by a process known as arthroconidiation [16]. The single cells then exhibit polarised growth before maintaining their shape and dividing by fission, at which point they are considered to be yeast cells. The specific host cell cues which drive direct *T. marneffei* morphogenesis are yet to be identified, and simple amenable hosts should be useful in facilitating such discoveries. Despite its public health significance, the range of infection models available for studying the disease caused by *T. marneffei*, is still highly limited compared to other fungal pathogens.

*Tenebrio molitor* larvae (mealworms) have previously been used to study a limited but diverse variety of human pathogens: Candida species (*C. albicans* and *C. tropicalis*), *Fonsecaea species* (*F. pedrosoi and F. monophora*) and *A. fumigatus* [19–22]. It has also been used for studying entomopathogenic fungi (*Metarhizium anisopliae and Beauveria bassiana*)[23–25], a fungal pathogen of plants (*Fonsecaea erecta*) [22], and the toxicity of various fungal metabolites [26]. Its tolerance to growth at 37°C makes this species particularly suitable for studying pathogens capable of infection establishment at human body temperature and is especially important for species that exhibit host- and/or temperature-specific developmental processes like *T. marneffei*. Further adding to this species’ utility for studies of fungal pathogenesis, is the growing number of research studies characterising *T. molitor* immune responses to invading microbes [27, 28]. Like other insects, *T. molitor* lack an adaptive immune system, but have a complex innate immune system with both cellular and humoral responses, similar to vertebrate innate immune systems [26]. As talaromycosis is prevalent in HIV-infected patients who have a dramatically compromised adaptive immune response, the lack of an innate immune system in *T. molitor* enhances their utility for modelling the opportunistic infection.

Here we describe a *T. molitor* model for talaromycosis that can be used routinely for testing and comparing the virulence of different *T. marneffei* mutant strains and/or clinical isolates. We have found that *T. molitor* larvae are susceptible to *T. marneffei* infection and that the larval host supports the production of *T. marneffei* yeast cells at 37°C, making *T. molitor* a suitable simple host for fungal virulence experiments. We also established a methodology for monitoring infection progression within larvae by microscopy, which can provide additional insights into the host-pathogen interaction that complements survival data.

Using this *T. molitor* infection model, we investigate the complex relationships between temperature, morphology and host cell damage by *T. marneffei*. Pathogenicity of *T. marneffei* is thought to be closely associated with its temperature-dependent yeast morphogenesis, however there are few opportunities to assess virulence of its hyphal form within host tissues. *T. molitor*, being ectothermic, presented a unique opportunity to assess morphogenesis and virulence within a host at 25°C. This model also allowed us to examine how directly inoculating pathogenic yeast cells impacts survival outcomes. Our results highlight the complexity of the human pathogen’s dimorphic switch and demonstrate distinct differences in the pathogenic potential of conidia, hyphae and yeast cells.

## Results

### Assessment of *T. molitor* thermotolerance

The optimum temperature range for *T. molitor* is 22-28°C [29], but their previous use in experiments at 37°C [19–22] suggests that the larvae can tolerate temperatures well beyond their optimum range. Before conducting a *T. marneffei* infection experiment, survival of two uninfected control groups was monitored over a 14-day period at 37°C to confirm suitability of the incubation conditions (i.e. temperatures, humidity and feed) and determine an appropriate duration for virulence experiments. Larvae in the first control group were not injected while larvae in the second group were injected with sterile Tween solution (vehicle-only injection control). All larvae were allowed to acclimate at 37°C for 16 hours prior to the inoculation/experiment ‘start’ time. Previously, El-Kamand et al. (2022) found that the survival of *T. molitor* larvae did not significantly differ when injected at different sternites, so we chose to inject the larvae at the third sternite as this improved safety by maximising the distance between the operator’s fingers and the syringe needle whilst maintaining control of the animal.

Survival was monitored for 14 days for three replicates. All larvae in the No-Injection control (NI control) group survived for the first 11 days post-injection (PI), and then 27% of the larvae died between days 11 and 14 (**Fig. S1A**). Survival of the Tween-injected control (Tween control) group was similar; with 21% of larvae dying 7-14 days PI. The cause of the late-stage deaths is unknown. Although the injection treatment was associated with a slightly earlier commencement of this late-stage decline, this decline occurred in both groups so could not be attributed solely to the injection treatment.

Survival of NI and Tween control larvae was also monitored for 14 days at 25°C. Survival of both groups was again mostly stable during the initial phase of the before beginning to decline (**Fig. S1B**). The survival decline did commence later in this experiment compared to the 37°C assay. At 25°C, persistent survival declines commenced at 9- and 12-days PI for the Tween and NI control groups respectively. It was also noted that some larvae began to pupate 13-14 days PI when incubated at 25°C, which did not occur at 37°C. These results indicated that larval survival (and pupation) was influenced, in part, by temperature and whether they had received an injection (primarily at 25°C), but was sufficiently stable such that survival assays could be conducted at either 25°C or 37°C. Due to the observed variability in control survival after 7 days, a 6-day monitoring period was selected for the subsequent *T. marneffei* virulence tests.

### Susceptibility of *T. molitor* to infection by *T. marneffei* at 37°C

To assess the susceptibility of *T. molitor* larvae to *T. marneffei* infection and determine a suitable dosage for virulence assays, larvae were injected with four different inocula of viable conidia (total number injected = 10^6^, 10^5^, 10^4^ and 10^3^) or sterile Tween solution. Larvae were monitored daily for survival (determined by responsiveness to touch) and changes in colour (production of melanin pigment). Larvae injected with 10^5^ or 10^6^ conidia showed statistically significantly reduced survival compared to the Tween-control group (**Fig. S2, Table 1**). Survival of the *T. marneffei*-infected larvae was dose-dependent, with larval survival decreasing with increasing inoculum. The group injected with 10 conidia showed a more consistent decline in survival over the 6-day infection period than the 10 group, which seemed to plateau at 4 days PI. Survival of larvae injected with 10 conidia was also more variable between replicate experiments, suggesting that this inoculum was near the threshold required for infection establishment. For this reason, the 10 inoculum was deemed most suitable for subsequent virulence screening.

**Table 1.**
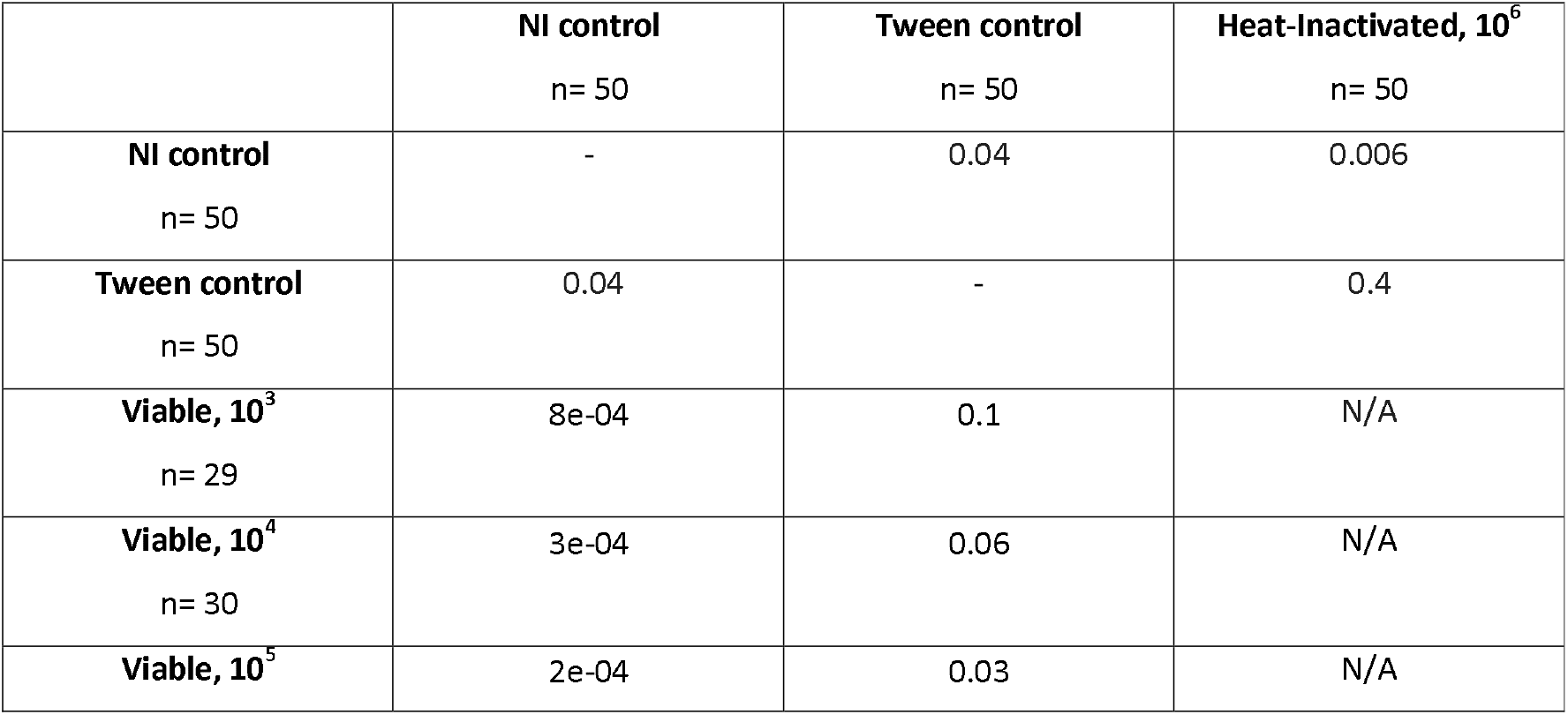
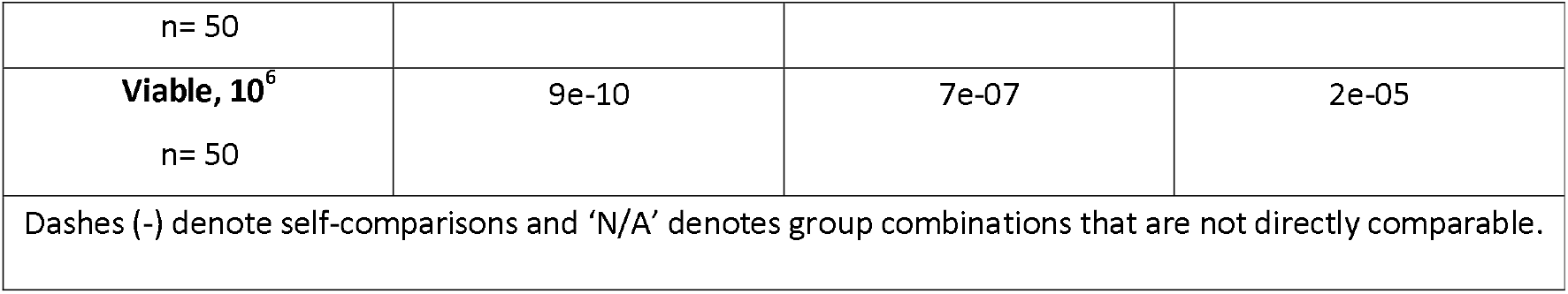
Pairwise LogRank test for differences in infected and non-infected *T. molitor* larvae at 37 °C.

To confirm that the observed deaths were attributable to the growth or activity of the live fungus rather than inherent toxicity of the injected conidia, survival of larvae injected with viable conidia was compared to those injected with the same number (10^6^) of heat-inactivated (HI) conidia. Survival of the HI group significantly differed from the viable conidia group but not from the Tween control group, indicating that survival outcomes were associated with conidial viability rather than the presence of conidia in the injected solution (**Fig. 1, Table 1**).

**Figure 1.**
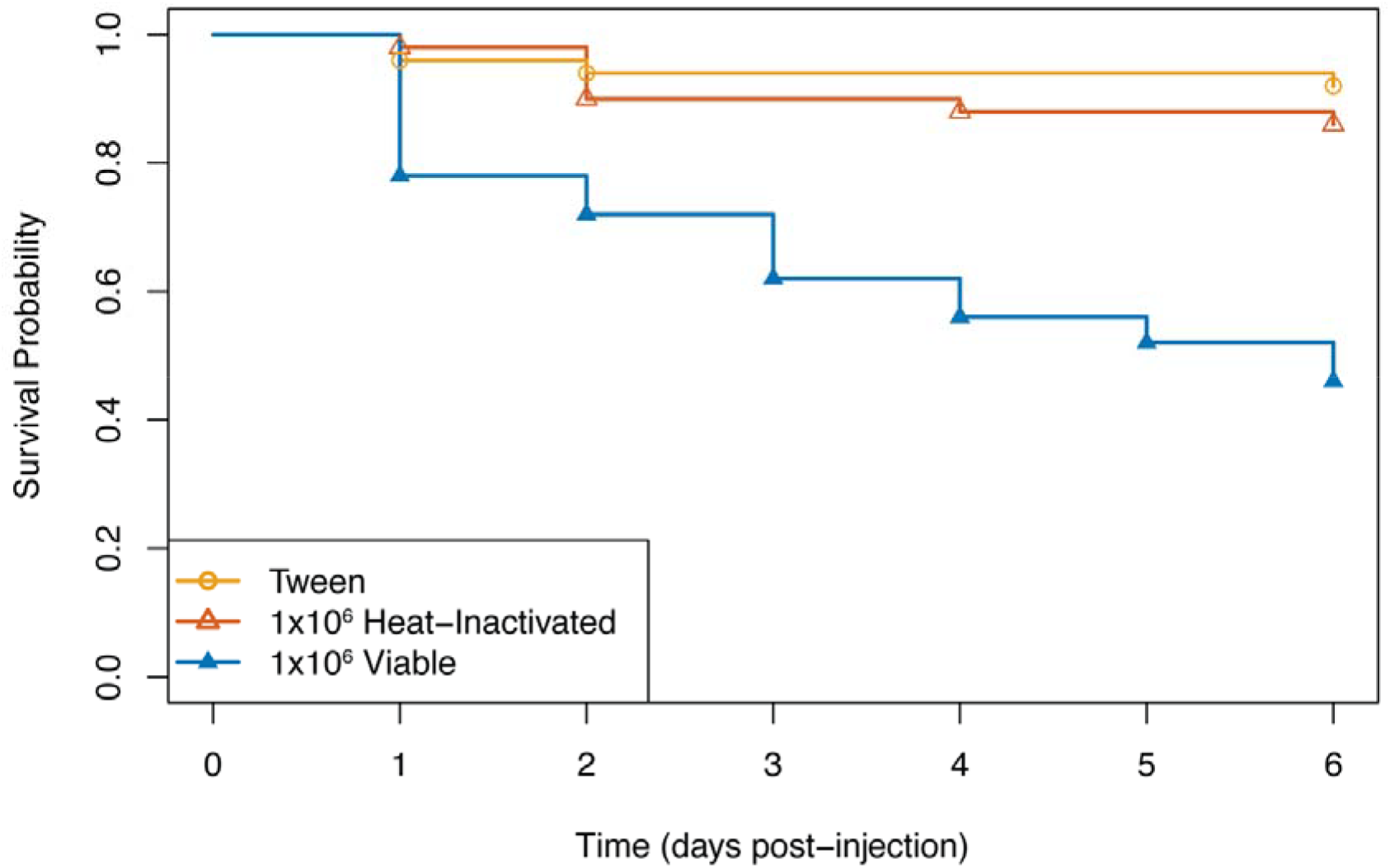
Survival of *T. molitor* larvae infected with *T. marneffei* at 37 °C. Kaplan-Meier plot showing survival probabilities for larvae injected with Tween solution alone or *T. marneffei* conidia in Tween solution over a 6-day period. Injection of larvae with 10^6^ viable conidia results in a significant reduction in survival compared to injection with Tween solution, while survival of larvae injected with 10^6^ heat-inactivated conidia does not significantly differ from those injected with the Tween solution (vehicle-only control) alone. Survival curves were built with pooled data from 5 replicate experiments (n= 50 per group).

24-48 hours before infected larvae succumbed to the infection, they were usually less active than uninfected control larvae; exhibiting more subdued responses to touch, such as only twitching their legs rather than crawling or moving their whole body. Many of the infected larvae also became increasingly melanised during the experiment, and 85.7% of dead larvae (including viable conidia, HI conidia and Tween-injected individuals) were melanised at the time that death was recorded, suggesting that melanisation is usually (although not always) indicative of poor larval health. It is noteworthy that while the presence of melanisation was strongly correlated with survival outcomes, the degree of melanisation was highly variable at the time of death, ranging from discrete patches of melanisation to whole-body discolouration (**Fig. S3**). Melanisation was also highly prevalent in larvae that had been injected with 10 conidia but were still alive 6 days PI (88.9% of surviving larvae injected with 10 conidia), suggested that their survival was more reflective of a delay in disease progression rather than a recovery (Table S1).

The infection model was established using the *T. marneffei* reference strain (FRR2161), and most molecular genetic studies use derivatives of the FRR2161 strain that are deficient in the non-homologous end joining (NHJEJ) pathway to facilitate genetic manipulation [30, 31]. As NHEJ-impaired strains are the recipient strain for most genetic studies, they are the most suitable ‘unmodified’ control for experiments assessing the virulence of genetically modified strains. To confirm that the infection model and associated conditions would also be suitable for such NHEJ-impaired laboratory strains and their derivatives, larvae were also infected with 10^6^ of the *niaD1ΔligD* strain (a routinely used NHEJ-deficient, nitrate reductase mutant). Virulence of the *niaD1ΔligD* strain did not significantly differ from that of the FRR2161 strain (**Fig. S4**), indicating that the infection conditions and 10^6^ inoculum should also be suitable for virulence experiments using this or comparable strains.

### Susceptibility of *T. molitor* to infection by *T. marneffei* at 25°C

To examine the relationship between *T. marneffei* virulence and temperature, larvae were injected with the same four inocula of viable conidia (total number injected = 10^6^, 10^5^, 10^4^ and 10^3^) or sterile Tween solution, incubated at 25°C and monitored daily for survival. Of the four concentrations of viable conidia tested, only the larvae injected with 10 conidia significantly differed from the Tween control group (**Fig S5, Table S1**). For the first 3 days, survival of larvae injected with 10^6^ conidia at 25°C declined at a similar rate to those that had been incubated at 37°C, however no additional deaths occurred for the 25°C group after 3 days PI (**Fig. 2**), suggesting that the remaining *T. marneffei*-infected larvae are recovering from the infection after this point. Despite this plateau at 25°C, the result of the log rank test comparing the two temperatures was not statistically significant (p-value = 0.2). Even with this non-significant result for larvae injected with 10^6^ viable conidia, it still appears that the development and progression of talaromycosis is less severe at 25°C than 37°C given that larvae require 10-fold higher doses of conidia before their survival is impacted.

**Figure 2.**
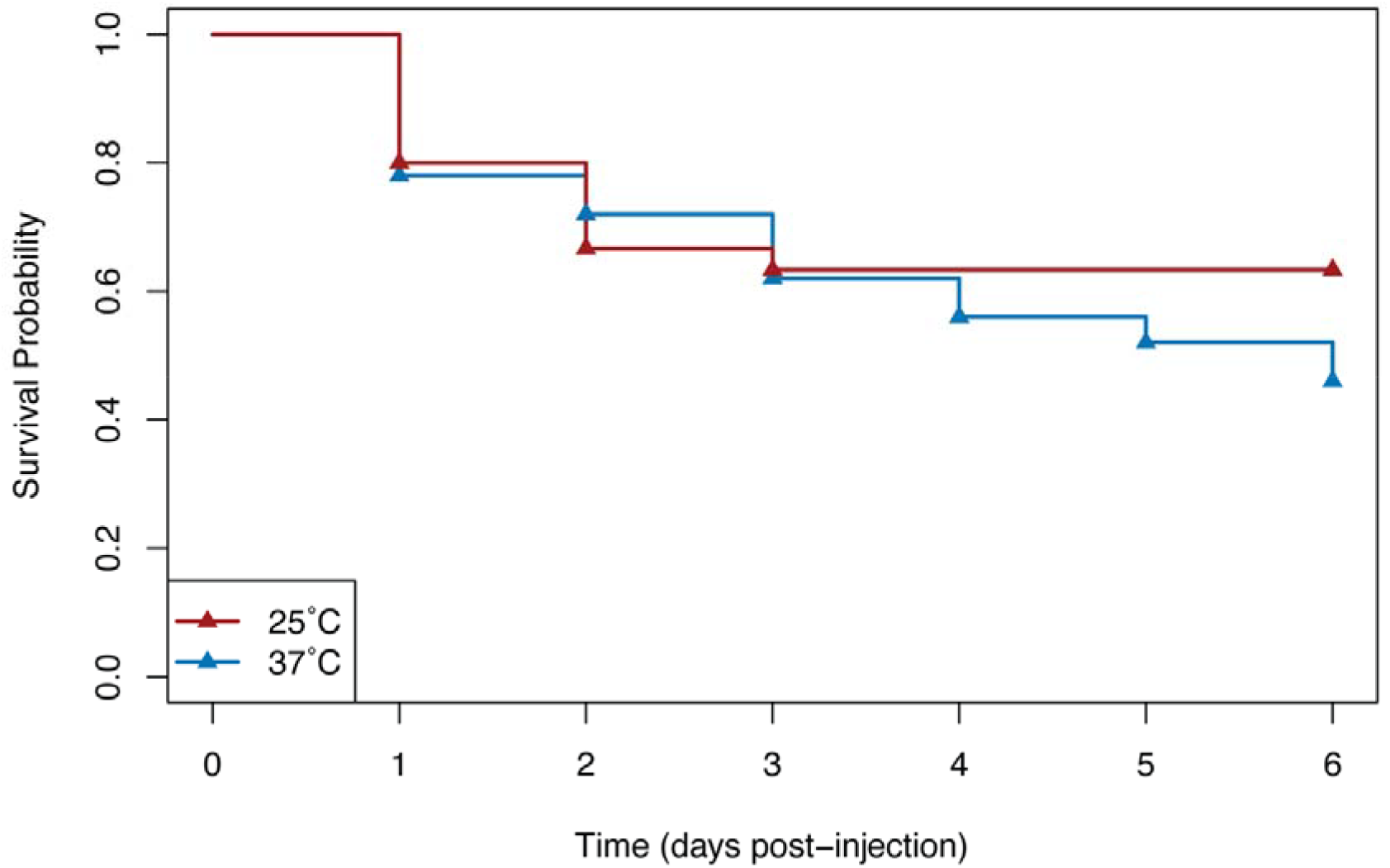
Survival of *T. marneffei*-injected larvae is temperature-dependent. Kaplan-Meier plot showing survival probabilities for larvae injected with 10^6^ conidia and incubated at either 25°C or 37 °C. Survival decline is similar for both temperatures for days 0-3 and then the two curves deviate. During days 3-6, survival continues to decline at 37 °C, but does not at 25 °C. The 25 °C and 37 °C survival curves were built with pooled data from three and five replicate experiments, respectively.

### Analysis of growth and morphogenesis within the *T. molitor* host

As *T. marneffei* is dimorphic with a yeast form that has only previously been observed within vertebrate hosts, we were also interested in determining which form of fungus was growing within this invertebrate larval host at each of the two temperatures. To examine *T. marneffei* morphology and distribution within the *T. molitor* host, *T. marneffei*-infected larvae were incubated at 25°C or 37°C and fixed at either 6- or 48-hours post-injection. NI control larvae were also incubated for 48 hours at each temperature and served as negative controls for this experiment. Whole *T. molitor* larvae were sagittally sectioned and stained with Hematoxylin and Eosin (H&E) to confirm integrity of the larval tissue following processing (**Fig. S6**), or Grocott’s Methenamine Silver (GMS) to stain for, and assess morphology and distribution of, *T. marneffei* within the larval section. At both 25°C and 37°C, the GMS-stained NI control sections were clear of silver staining (**Fig. S6**).

Stained *T. marneffei* conidia were identifiable within the tissues of infected larvae 6 hours post-injection at both 25°C and 37°C (**Fig. 3B**). No differences in size or shape were identified between the two temperatures, and many conidia were swollen, consistent with an initial period of isotropic growth before cell polarisation (production of a germ tube or yeast morphogenesis). Conidia were primarily confined to the anterior half of the larvae at this early time point (**Fig. 3A**). Overall, *T. marneffei* distribution and morphology was indistinguishable at 25°C and 37°C at 6 hours post-injection.

**Figure 3.**
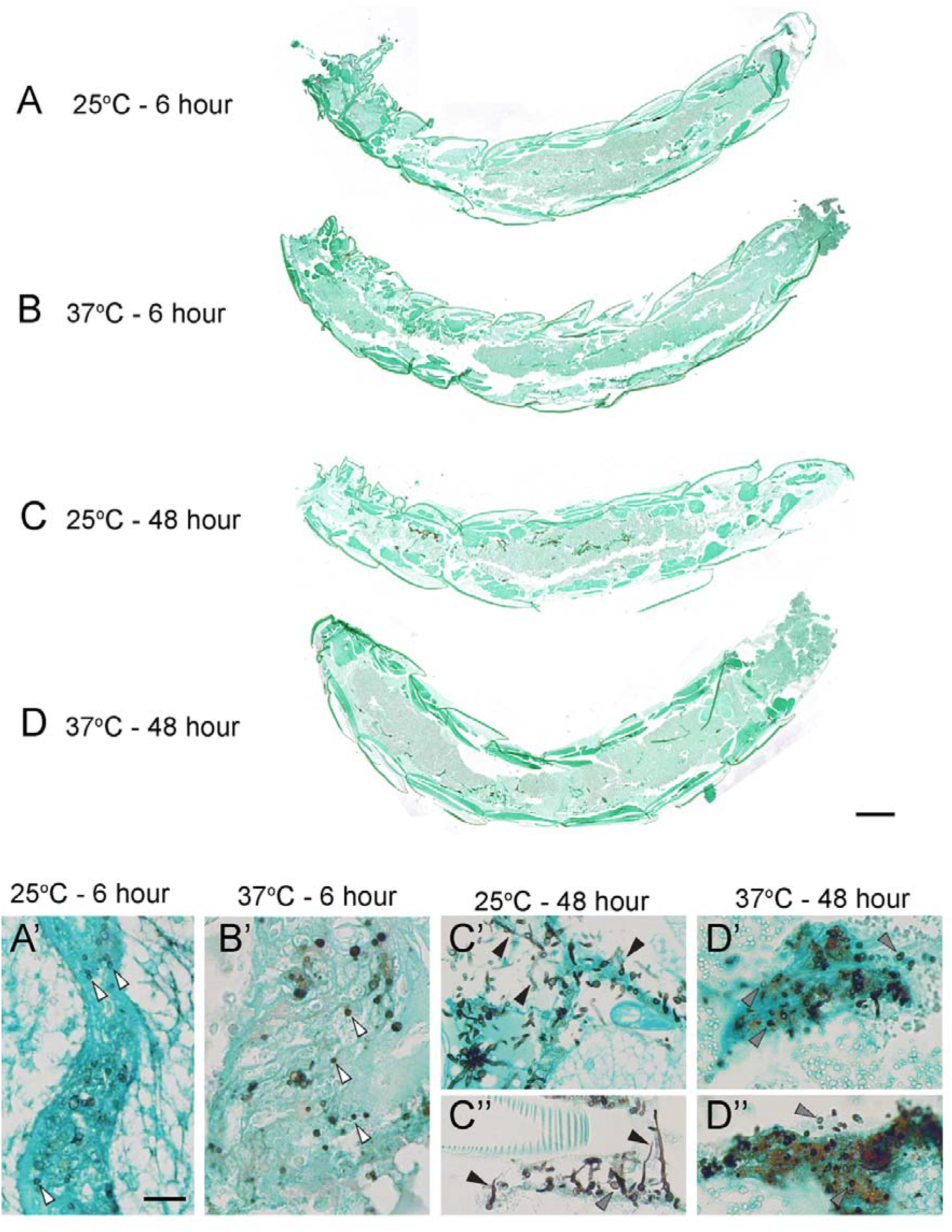
*T. marneffei* morphology within *T. molitor* host tissues. (A-D) GMS-stained sagittal sections of *T. marneffei*-infected larvae 6 (A, B) and 48 (C, D) hours post-injection (anterior end to the left, posterior end to the right) following incubation at 25°C (A, C) or 37°C (B, D). (A’-D’, C’’, D’’) Higher magnification images show examples of stained *T. marneffei* in each condition. (A’, B’) At 6 hours post injection, *T. marneffei* conidia (white arrowheads occurred in patches of relatively low density primarily as spores after incubation at both temperatures. (C’, C’’) After 48 hours post injection at 25°C, dense patches of fungal growth were observed across the entire length of *T. molitor*. Growth is composed primarily of hyphal filaments (black arrowheads), as well as some yeast cells (grey arrowheads). (D’, D’’) After 48 hours post injection at 37°C, patches of dense *T. marneffei* were found across the entire length of *T. molitor*, with the vast majority exhibiting round yeast morphology (grey arrowheads). Brown discolouration was also apparent in the tissue surrounding the yeast cell and not observed in the other conditions. Scale bar = 1 mm in D (for A-D) and scale bar = 20 μm in A’ (A’-D’, C’’, D’’).

The distribution of *T. marneffei* was more uniform 48 hours post-injection at both temperatures, with dense patches of GMS-stained fungal growth observed across the entire anterior to posterior axis. Morphological differences were, however, clearly apparent at the later timepoint. At 37°C, ovoid yeast cells were the dominant cell type observed (**Fig. 3C**), indicating that the *T. molitor* host environment could support yeast cell development. Some hyphae were also identified, however these make up < 5% of the fungal growth at 37°C. Interestingly, many of the yeast cells were present in dense clusters and overlapping each other, as though they were confined within discrete areas of host tissue. These tissue regions also contained large amounts of brown pigmentation. In contrast, primarily hyphal growth was detected in the *T. marneffei*-injected larvae at 25°C (**Fig. 3D**), consistent with temperature being a major regulator of dimorphic switching. However, yeast-shaped cells were also present at the lower temperature (approximately 10-20% of total fungal growth). This combination of morphologies is similar to 37°C *in vitro* cultures, which are typically a mixture of hyphae, arthroconidia and yeast cells [32]. It was also noted that the hyphal growth at 25°C was less spatially confined than the clusters of yeast cells at 37°C and associated with less brown pigmentation.

Surprisingly, more fungal material was detected overall in the larval sections at 25°C than at 37°C after 48 hours. This was unexpected considering that larval susceptibility to *T. marneffei* infection was lower at 25°C in the survival experiments (evidenced by lower overall deaths when larvae were injected with the same numbers of conidia). As haemolymph cannot be fixed, it is possible that fungal cells that had not adhered to larval tissues were lost during sample processing. To determine whether fungal material was also present in the circulating haemolymph, haemolymph samples were collected from infected larvae following incubation at either 25°C or 37°C for either 6 or 48 hours, as well as from Tween and NI control larvae incubated at 25°C or 37°C for 48 hours, and stained with calcofluor white.

Insect cells of 5-20 μm in length were visible under differential interference contrast (DIC) microscopy in all haemolymph samples collected, consistent with previously documented sizes of prohemocytes, granulocytes and plasmocytes [33, 34]. Cell morphology ranged from spherical (likely prohemocytes and granulocytes) to more elongated (likely plasmocytes), and were mostly found in large multi-cell clusters, such that it was difficult to discern cell borders clearly (**Fig. S7**). Calcofluor-stained conidia (2-3 μm in diameter) were observed in the haemolymph samples taken 6 hours post-injection at both 25°C and 37°C (**Fig. S8**). The conidia were associated with insect haemocytes, but sparsely distributed, such the majority of haemocytes observed were not associated with conidia. In the 37°C/48-hour samples, small clusters of yeast cells (4-6 μm in length) were consistently observed, including cells that were dividing at the time of fixation (**Fig. 4**). The yeast cells were observed more frequently than the conidia, and often in groups of dividing cells suggestive of proliferation from a single conidium. These yeast cells were always observed in association with *T. molitor* haemocyte clusters and appeared to be spatially confined to a small area, possibly indicative of an encapsulation response. In contrast, almost no fungal material was detected in 25°C/48-hour sample (**Fig. S9**), with the exception of one small group of yeast-like cells (**Fig. S10**). This strongly suggests that the hyphal form of the fungus which is dominant at 25°C degrees, remains adherent to the insect tissues, rather than circulating within the haemolymph. Fungal growth was not observed in any of the control haemolymph samples (NI and Tween controls incubated at 25°C or 37°C) and no differences were observed between the four control conditions as expected (**Fig. S9**). This haemolymph data shows fungal staining of histological sections is informative for the morphology and distribution of infecting fungus, but does not represent the total fungal burden.

**Figure 4.**
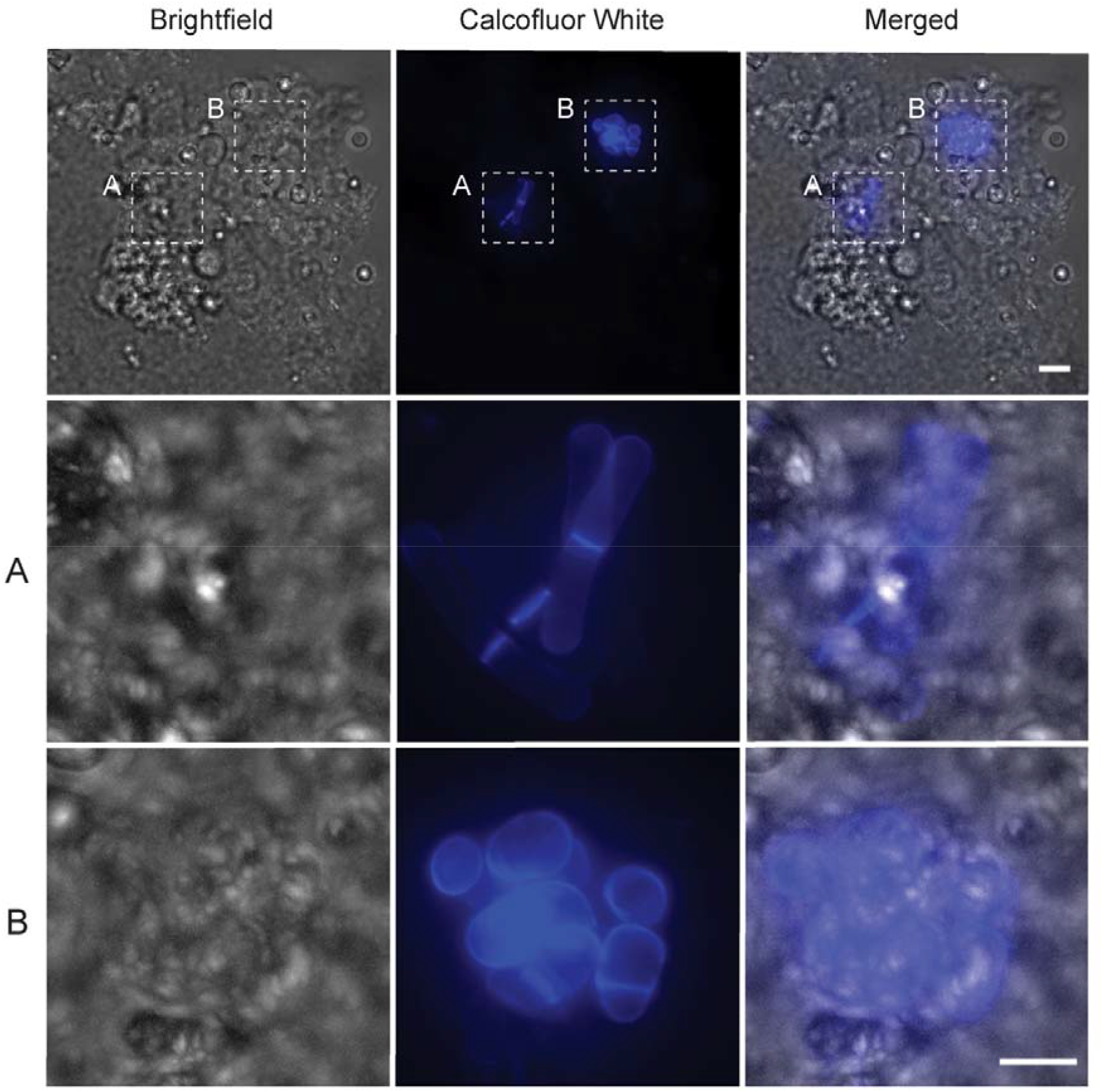
Yeast cells present in haemolymph 48 hours post-injection and incubation at 37°C. Micrograph of *T. molitor* haemolymph collected from larvae incubated at 37°C for 48-hours post-injection with 10^6^ *T. marneffei* conidia and stained with 10 μg/ml calcofluor white to observe fungal cells. Two discrete clusters of yeast cells (stained with calcofluor white) are associated with a cluster of haemocytes (visible under brightfield). Stained septa within the yeast cells are consistent with active replication of yeast cells at the time of fixation. The two groups appear to be spatially confined to small regions of clustered haemocytes, potentially indicative of encapsulation by the haemocytes (Merged). Scale bar for top row is 10 μm, and scale bar for inset panels (second and third rows) is 5 μm.

### Yeast cell virulence

To investigate whether the differences in survival between larvae infected at 25°C and 37°C could be explained by yeast cells having a greater capacity for host damage, we examined whether injection with yeast cells instead of conidia further reduced survival. THP-1 human macrophages were infected with *T. marneffei* conidia and after 24 hours the macrophages were lysed and filtered to generate a cellular suspension comprised primarily of ex *vivo* yeast cells (approximately 2/3 yeast cells and 1/3 short hyphal fragments). Viability of the yeast cell and conidial suspensions was verified on Synthetic Dextrose medium (Table S1).

Larvae were injected with either 10^6^ cells of either conidia or ex *vivo* yeast (total number of cells including hyphal fragments) and monitored daily for survival. As yeast cells were suspended in phosphate buffered saline (PBS) instead of the Tween solution (the vehicle solution for the conidia), a PBS-only control was used in the assay. Survival of larvae injected with PBS did not differ from those injected with the Tween solution (p-value = 0.4), confirming that the alternative diluent does not influence survival. Larval survival was significantly reduced when injected with the yeast cell suspension compared to injection with the same number of conidia (**Fig. 5, Table 3**). 60% of the conidia-injected and 87% of the yeast-injected larvae had died by the conclusion of the experiment. As in the previous 37°C survival assay, most of the surviving infected larvae (conidia and yeast-injected) were melanised (75%) and/or slow to respond at the end of the 6-day monitoring period, suggesting they may have succumbed to infection in the coming days had the experiment continued. The same yeast versus conidia experiment was also performed with *niaD1ΔligD* strain with similar results (**Fig. S11, Table S2**). Survival of larvae injected with *niaD1ΔligD* yeast cells did not significantly differ from those injected with FRR2161 yeast (p-value = 0.3). Based on the survival data and these observations, we conclude that yeast cells have a greater capacity for host damage than conidia, and that direct injection of yeast cells leads to a quicker establishment of disease, most likely due to bypassing the period required for morphogenesis of yeast cells from conidia.

**Table 2.** Pairwise LogRank test for differences in infected and non-infected *T. molitor* larvae at 25 °C.

|  |  |  |  |
| --- | --- | --- | --- |
|  | **No Infection control**<br>n= 30 | **Tween control**<br>n= 30 | **Heat-Inactivated, $10^6$**<br>n= 30 |
| **No Infection control**<br>n= 30 | - | 0.04 | 0.08 |
| **Tween control**<br>n= 30 | 0.04 | - | 0.7 |
| **Viable, $10^3$**<br>n= 30 | 0.2 | 0.4 | N/A |
| **Viable, $10^4$**<br>n= 30 | 0.08 | 0.7 | N/A |
| **Viable, $10^5$**<br>n= 30 | 0.08 | 0.7 | N/A |
| **Viable, $10^6$**<br>n= 30 | 3e-04 | 0.04 | 0.01 |
| Dashes (-) denote self-comparisons and 'N/A' denotes group combinations that are not directly comparable. |  |  |  |

**Table 3.** Pairwise LogRank test for differences in larval survival when injected with conidia or yeast cells.

|  |  |  |  |
| --- | --- | --- | --- |
| | Viable Conidia, $10^6$<br>n= 30 | PBS control<br>n= 30 | Tween control<br>n= 30 |
| PBS control<br>n= 30 | N/A | - | 0.4 |
| Viable Conidia, $10^6$<br>n= 30 | - | N/A | 3e-05 |
| Viable Yeast cells, $10^6$<br>n= 30 | 4e-04 | 5e-10 | N/A |
| Dashes (-) denote self-comparisons and 'N/A' denotes group combinations that are not directly comparable. |  |  |  |

**Figure 5.**
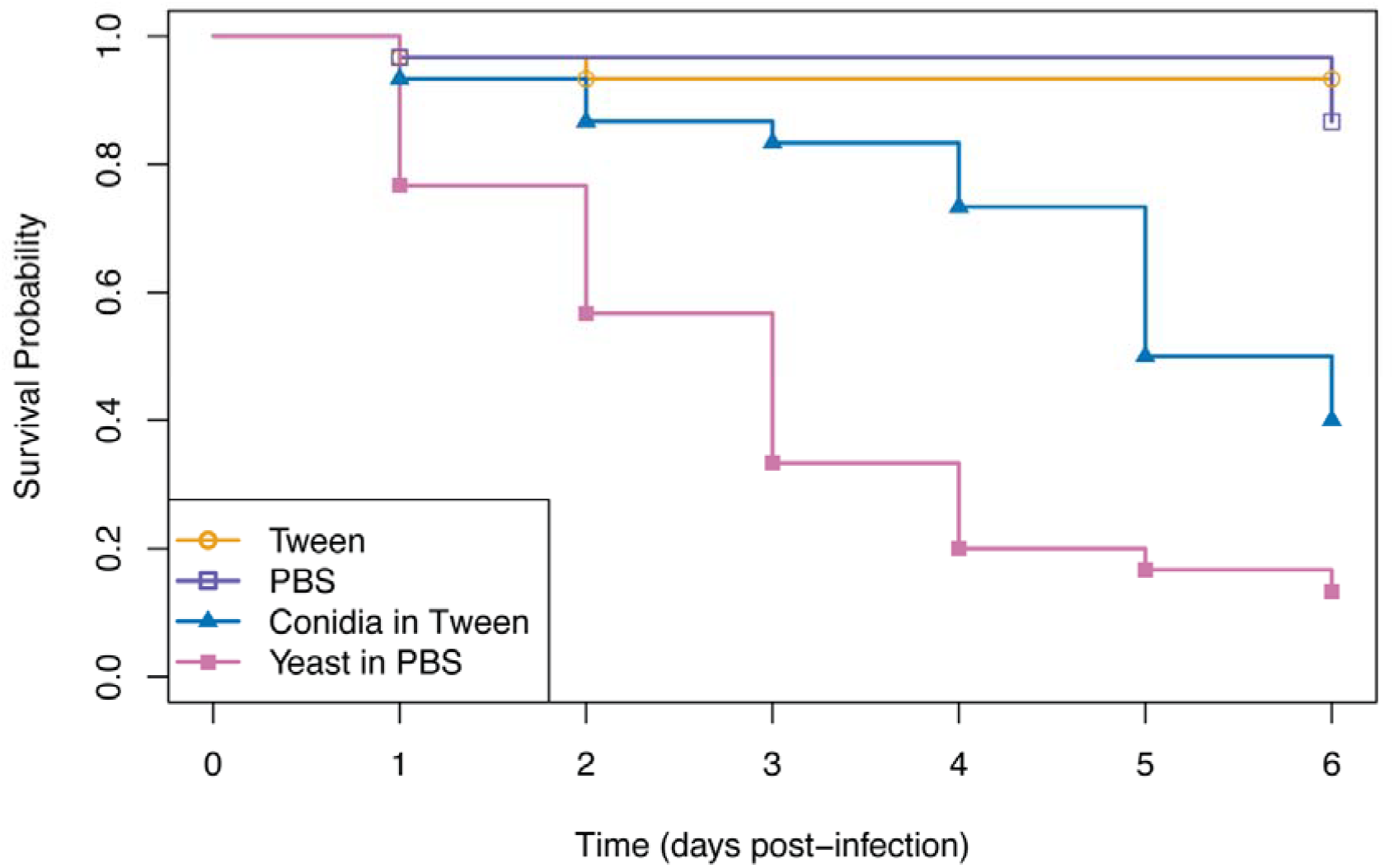
Differences in survival of larvae infected with 10^**6**^ conidia and ex *vivo* yeast cells 37 °C. Kaplan-Meier plot showing survival of larvae injected with viable conidia in Tween, viable ex *vivo* yeast cells in PBS, Tween solution or PBS (vehicle only controls) and incubated at 37 °C. Direct injection with *T. marneffei* yeast cells increased the rate of survival decline compared to injection with *T. marneffei* conidia. There was no significant difference in survival of larvae injected with Tween solution or PBS, confirming that neither vehicle solution impacted survival outcomes. Survival curves were built with pooled data from three replicate experiments (n= 30 per group).

## Discussion

Host-pathogen interaction models are critical for investigating fungal pathogenesis, and animal infection models offer especially important insights into the development and progression of fungal infections within a whole animal – something that is not possible in cell-based infection experiments. A small but diverse range of models have previously been developed to examine *T. marneffei* pathogenesis within a whole animal, and each come with their own advantages and disadvantages. Adding to the toolkit for this species, we have developed the first *T. molitor* model for *T. marneffei* infections – a simple, thermotolerant host that can be used to study *T. marneffei* dimorphism and pathogenesis in any lab already certified for work with genetically-modified *T. marneffei*.

Three other invertebrate hosts have been used for studying *T. marneffei* previously: nematodes (*Caenorhabditis elegans*), fruit flies (*Drosophila melanogaster*), and wax moth larvae (*Galleria mellonella*) [35–40]. Despite being ectothermic, nematodes and fruit flies both have temperature ceilings below 37°C, so pathogenicity was only investigated at 25°C and 29°C respectively in these species. This temperature restriction is an issue for studying thermally dimorphic pathogens like *T. marneffei*, as yeast cell development requires temperatures of at least 32°C, and only occurs optimally at 37°C [41]. The *T. marneffei* transcriptome also changes significantly upon exposure to 37°C [42, 43], such that many genes (including those implicated in virulence) may not be appropriately expressed in these infection models. The other invertebrate model, *G. mellonella*, is most similar to the *T. molitor* model, as it is also tolerant to growth at 37°C. *G. mellonella larvae* are available in several countries and are the most widely used model for human fungal pathogens due to their thermotolerance. This species is not, however, readily available in all countries (such as Australia and Brazil), therefore requiring researchers to invest time and resources if they are to maintain their own colony [44]. In contrast, *T. molitor* larvae are more broadly available and commonly sold as animal feed, permitting researchers to order larvae for infection experiments when required.

In this study, four different inocula of the *T. marneffei* reference strain (1 × 10^3^, 1 × 10^4^, 1 × 10^5^ and 1 × 10^6^) were evaluated in *T. molitor* larvae at 37°C to assess their susceptibility to *T. marneffei* infection and determine an appropriate concentration of conidia for fungal virulence assays. It was found that injection of ≥ 1 × 10^5^ conidia significantly decreased survival compared to injecting with the Tween solution alone at 37°C, although 1 × 10^6^ conidia was deemed to be the most suitable for *T. marneffei* virulence screening as it produced the most consistent survival decline across a 6-day monitoring period. This is equivalent to the concentrations of *T. marneffei* conidia that have been used in *G. mellonella* infection experiments at 37°C, where 1 × 10^5^ conidia was also the lowest dosage to significantly impact larval survival compared to control injection [38], and inocula of 1 × 10^6^ or 1 × 10^7^ conidia were subsequently used to assess virulence of different *T. marneffei* strains [39, 40]. Direct comparisons between these *G. mellonella* infection experiments and those described here for *T. molitor* are complicated by the fact that the *G. mellonella* studies used various clinical or ‘wild type’ isolates of *T. marneffei* rather than the FRR2161 reference strain, and none of these strains were available to test. The FRR2161 reference strain is the progenitor of key laboratory strains such as *niaD1ΔligD*, which is defective in non-homologous end joining repair to facilitate efficient genetic modification [30]. The *niaD1ΔligD* strain was also assessed as part of this study, as this strain and genetically modified derivative strains are likely to be tested for virulence with this model. Our findings indicate that 1 × 10^6^ conidia is also a suitable inoculum when screening virulence of NHEJ-impaired strains like *niaD1ΔligD*.

Dosages used in laboratory disease models are often greater than is necessary to cause serious disease in the host being studied [45]. To this end we aimed to initiate infection with as few conidia as possible that provided a robust and reproducible effect on survival. This chosen concentration was low enough that all larvae were not killed within the 6-day experimental period, providing scope to detect both reductions and increases in virulence. As shown to be in the case in each of the other invertebrate models [35, 36, 40], it is highly likely that *T. marneffei* virulence will vary between strains of different origins such as clinical isolates or laboratory produced mutants. Furthermore, some control larvae (Tween or NI) survived well past the 6-day monitoring period that was initially selected, suggesting that infected larvae could be monitored for a longer period if required (e.g. if examining delayed virulence phenotypes or isolates associated with slower disease progression).

This is the first study of *T. molitor* (or any invertebrate model for *T. marneffei* infection) where whole animal histology has been performed. By fixing and sectioning whole larvae we were able to examine not only the morphology of this pathogen, but also its distribution within a host. Doing this at both 6 and 48 hours also provided insight into infection development, showing a shift from anterior-localised conidia to yeast cells that were distributed throughout the larva. The detection of *T. marneffei* yeast cells within larvae after 48 hours is important for two key reasons. Firstly, because this is the cell type that is found in infected human hosts [13], meaning that virulence screens using this model are assessing the human disease-causing form of the fungus. The second reason is that it provides information about the environmental cues that induce development of the pathogenic cell type. In the absence of host cues, such as at 37°C *in vitro*, cultures are composed of highly branched hyphae and yeast cells produced by arthroconidiation. These yeast cells are usually more elongated than those that develop within mammalian macrophages, which are more oval shaped [46]. The presence of ovoid-yeast cells and the absence of hyphal fragments at 48 hours PI at 37°C indicates that the *T. molitor* host environment is conducive to yeast cell morphogenesis. This finding is consistent with yeast cells being found in the tissues and hemolymph of *G. mellonella larvae* [39, 40], and suggests that the host factors that stimulate this direct yeast cell germination are conserved in invertebrates.

Due to the observed clustering of haemocytes in the haemolymph analysis, we were unable to conclusively determine whether the *T. marneffei* yeast cells identified were developing intracellularly. However, based on what is known about the sizes and functions of different *T. molitor* haemocytes, it seems most likely that *T. marneffei* conidia and yeast cells are encapsulated rather than phagocytosed. Granulocytes, the only haemocyte type known to phagocytose invading pathogens, are typically ~10 μm in diameter [33]. Therefore, these granulocytes could contain 1-2 yeast cells at most, and are too small to physically contain the larger clusters of yeast (up to 10 yeast cells) that were observed in close association with one another in the haemolymph samples. Instead, aggregation of haemocytes around the fungal cells would explain why the *T. marneffei* yeast cells still appeared to spatially confined by the haemocytes they were associated with. Plasmocytes, which represent 23-28% of all haemocytes, are deemed the cell type most likely responsible for encapsulation in *T. molitor*. They are known to be far more polymorphic but are often larger and more elongated than granulocytes (estimated to range from 9-15 μm in length) [34]. Cells exceeding a length of 10 μm were detected in the analysis, suggesting that this encapsulating cell type was present within the haemolymph analysed. Encapsulation has also previously been reported to be a primary immune reaction of *G. mellonella* to *C. neoformans* [47], and could potentially be the key cellular mechanism by which insects defend themselves against larger microbes, perhaps reserving phagocytosis for smaller microbes like bacteria. The hypothesis that *T. marneffei* is not phagocytosed in *T. molitor* raises questions about what host signals are driving yeast cell development at 37°C. Despite the fact that *T. marneffei* yeast are readily found within vertebrate macrophages, it is unlikely that the act of phagocytosis is the sole driver of yeast morphology, as homogenous yeast cell development is observed within macrophages but not within other phagocytic cells such as neutrophils in a 33°C zebrafish model [17]. More probable is that the larval hemolymph contain compounds similar to that of the macrophage phagolysosome.

It should also be noted that the large amounts of brown pigmentation were observed in close proximity to the yeast cells in the tissue sections, which may represent an accumulation of melanin. Melanogenesis, which is the production of brown-black pigmentation, commonly accompanies both encapsulation and phagocytic responses in arthropods [48]. It is hypothesised that intermediates produced during the production of melanin facilitate killing of invading microorganisms through the production of cytotoxic intermediates, whilst the final melanin product helps to protect the insect tissues from this cytotoxic damage by sequestering free radicals [48].

While the findings from the conidia-injection experiments at 37°C are most relevant for human disease, specific examination of virulence of the hyphal form and the yeast form provided greater insight into the capacity that the different fungal morphologies have for host damage. Direct injection of larvae with yeast cells isolated from infected macrophages resulted in the greatest number of total deaths, followed by conidia at 37°C (95% yeast cells after 48 hours), and then conidia at 25°C (80-90% hyphae after 48 hours). The difference in total deaths between injection with yeast cells and conidia at 37°C likely reflects the delay in producing yeast cells when using conidia, which must first germinate. This suggests that yeast cells are the most damaging for the host, which has been long-been considered the pathogenic form of the fungus but never directly proven. The difference in total deaths between injection with conidia and yeast cells at 37°C most likely reflects the additional germination period before conidia germinate into the damaging yeast cells.

Modulating the incubation temperature of the larvae during infection was particularly useful for investigating the influence of temperature on *T. marneffei* virulence and morphology in a host context – something that is not possible with live endothermic hosts or mammalian cell lines. Overall susceptibility of *T. molitor* to *T. marneffei* was reduced at 25°C, requiring 10 times more conidia to cause significant disease than at 37°C. This is despite the fact that infected larvae were still extensively colonised at the lower temperature. Whilst *T. molitor* may have improved anti-microbial immunity at 25°C, this does not seem to inhibit hyphal growth, nor the production of some yeast-like cells, within larval tissues. This supports the notion that yeast cells may secrete factors that are more damaging to the host than hyphae, and that the presence of fungal growth is not the primary driver of morbidity and mortality. It also suggests that the number of yeast cells detected in the 48hr/25°C tissue sections was too few to cause a sustained infection. As histology was not performed for larvae more than 48 hours PI, it is not known whether the survival plateau observed at 25°C (3 days PI) is associated with an overall reduction of fungal load within the tissues.

With this ectothermic host we have been able to study *T. marneffei* morphology and the associated impact on pathogenicity at both environmental and human body temperatures. We find that both of these traits are intricately linked, but that host signals also drive production of the most-damaging cell type in the absence of the temperature signal. As understanding of *T. molitor* immunology and genetics continues to accumulate, it may soon become possible to manipulate or disrupt specific immunity-related genes or signalling pathways to interrogate the fungus’ sensitivity to different conserved host responses, making this an invertebrate model especially useful for studying this, and other, dimorphic fungal pathogen of humans.

## Materials & Methods

### Fungal strains and growth conditions

The *T. marneffei* reference strain (FRR2161) and its derivative *niaD1ΔligD* strain (G821) were used in this study. FRR2161 was isolated from *Rhizomys sinensis* (bamboo rat) (supplied by J. Pitt, CSIRO Food Industries, Sydney, Australia). Conidial suspensions were prepared in sterile 0.005% v/v Tween 80 in dH_2_0 as previously described [32]. Conidial concentrations were determined using a haemocytometer and the suspensions serially diluted in sterile Tween solution. Conidia were heat-inactivated by incubation at 65°C for 30 min. To assess viability of the conidial suspensions, 10 μl of each suspension was plated on Synthetic Dextrose (SD) medium supplemented with 10 mM ammonium sulphate and incubated at 37°C for 6 days; growth of the viable conidia was proportional to concentration and no growth was observed for the heat-inactivated conidia.

To prepare suspensions of *T. marneffei* in *vivo*-derived yeast cells, THP-1 macrophages were infected with *T. marneffei* conidia. THP-1 cells were cultured in RPMI medium containing 10% foetal bovine serum at 37°C with 5% CO_2_. Cells were counted in a haemocytometer and 1×10^7^ cells were seeded into one 75 cm^2^ flask. Phorbol myrisate acetate was added to a final concentration of 32 nM and the cells incubated for 24h to induce macrophage differentiation. Unadhered THP-1 cells were removed by discarding the medium and fresh RPMI medium added, to which 1×10 *T. marneffei* conidia were seeded. THP-1/*T. marneffei* were co-incubated for 2h to allow phagocytosis. Unphagocytosed *T. marneffei* conidia were removed by discarding the medium and gently washing the adhered THP-1 monolayer with three 10ml changes of pre-warmed phosphate-buffered saline (PBS). Fresh RPMI medium was then added, and cells were incubated at 37°C for 24h. The medium containing detached THP-1 cells and *T. marneffei* growing outside of THP-1 cells was discarded. To extract *T. marneffei* yeast cells, 10 ml of ice-cold PBS was added and the infected THP-1 cells were detached by scraping with a sterile plastic spreader. To lyse the THP-1 cells the suspension was transferred to a centrifuge tube and drawn up into a syringe with a 25G needle and expelled 3 times. Microscopic examination showed that almost all THP-1 cells were ruptured releasing the *T. marneffei* yeast cells. Yeast cells were pelleted by centrifugation at 4000 g for 5 min, resuspended in 500 μl of PBS and counted in a haemocytometer. The cell suspension was then either diluted or pelleted again and resuspended in smaller volume of PBS to a final concentration of 1×10^8^ cells/ml. The conidia and yeast cell suspensions used in the yeast versus conidia infection assays were diluted and plated for viability (Table S3) on the same day as larvae were injected to ensure that viable counts were representative of the viability at the time of injection.

### Maintenance and infection of *T. molitor* larvae

*T. molitor* larvae were purchased from two stockists: BioSupplies (Yagoona, New South Wales) and Minibeasts Enterprises (Bannockburn, Victoria). The virulence assay was based on two previous studies [19, 20]. *T. molitor* larvae were size selected (150-200 mg) and checked for uniform colour and response to stimuli. Larvae were ‘housed’ in Petri dishes (10 larvae per dish) containing approximately 4 g of rearing diet and a 500 mg piece of frozen carrot. Rearing diet comprised of a 5:1 w/w ratio of wheat bran and LSA mix (linseeds, sunflower seeds and almonds). The larvae were incubated at the appropriate temperature (25°C or 37°C) overnight to allow them to recover from transport and acclimate to the incubation temperature prior to infection. Prior to injection, larvae were chilled to 4°C for 10-15 minutes to limit their movement and reduce the risk of injection injury. Groups of larvae were injected at the third sternite with an injection volume of 10 μl using a BD Ultra-Fine Insulin Syringe (0.33 mm × 12.7 mm, 29G) containing sterile Tween solution, sterile PBS, conidial suspension (diluted in Tween solution), or yeast cell suspension (diluted in PBS). Larvae were then returned to the appropriate incubator (25°C or 37°C). The ‘No Injection’ control group were also chilled at 4°C before being returned to the incubator but were not injected with any solution. Larval survival was then monitored every 24 hours for 6-14 days. No Injection control groups were included in all experiments to verify the health of the different larval batches. Survival was determined based on whether the larvae responded to physical stimulation. The piece of frozen carrot was replaced daily during the survival checks, and every 3-4 days the larvae were transferred to a new Petri dish with fresh rearing food.

### Survival data analysis

Kaplan-Meier survival analyses were performed using the Survival Analysis package in R. [49, 50] (https://CRAN.R-project.org/package=survival). Survival curves were generated using pooled data from at least three biological replicates. Larvae were censored either at the final observation time point if death had not occurred, or earlier if monitoring ended before the experiment endpoint. Survival curves were compared in a pairwise manner using the log-rank test and statistical significance was defined as p <0.05.

### Histology of *T. marneffei*-infected larvae

Larvae that had not died within the designated incubation period (6 or 48 hours) were selected for histological analysis and chilled to 4°C. As the larval exoskeleton is resistant to known fixative reagents, the first and last segments were removed from the pre-chilled larvae using a scalpel. Each larva was fixed in Davidson’s solution (2:3:1:3 ratio of 37% formaldehyde : absolute ethanol : acetic acid : dH_2_O) for 1 hour at room temperature and then 10% neutral buffered formalin (NBF) for 5 days at room temperature. Larvae were then transferred into an embedding cassette and processed in a Sakura Tissue-Tek VIP6 using the following cycling conditions: 10% v/v NBF for 2 hours, 50% v/v ethanol for 3 hours, 90% v/v ethanol for 3 hours, 3 × cycles of absolute ethanol for 3 hours, ethanol-xylene solution (50% v/v ethanol 50% v/v xylene) for 2 hours, 2 × cycles of xylene for 3.5 hours and 4 × 3 hour cycles of paraffin embedding at 58°C. The paraffin-embedded larvae were then sectioned sagittally (6 μm sections) using a Microm HM 325 microtome (Thermo Scientific) and paraffin sections were mounted onto Superfrost Plus microscope slides (Epredia).

Before staining, slides were dewaxed in xylene and hydrated through graded ethanol/dH_2_O (absolute, 90% v/v, 70% v/v, dH_2_O). Hematoxylin and eosin (H&E) staining was performed as per standard methods [51]. Silver staining was performed as follows: slides were oxidised in 5% w/v chromic acid for 1 hour. Slides were then washed with running tap water for 5 minutes, rinsed with dH_2_O and residual chromic acid was removed using 1% w/v sodium bisulphite for 1 minute. Slides were then washed in running tap water and rinsed thoroughly in dH_2_O, before being incubated in Grocott’s Methenamine Silver (GMS) solution (0.023% w/v silver nitrate, 1.44% v/v methenamine and 0.19% w/v sodium tetraborate in dH_2_O) at 60°C until positive control sections appeared golden brown with dark brown-black fungi (approximately 60 minutes). Slides were then thoroughly rinsed in dH_2_O, toned in 0.1% w/v gold chloride for 2 minutes and rinsed in dH_2_O again. Unreduced silver was removed using 2% w/v sodium thiosulphate for 2 minutes and slides were washed with running tap water for 5 minutes. Slides were counterstained with light green solution (0.04% w/v light green SF and 0.04% v/v acetic acid in dH_2_O) for 30 seconds. Stained slides (GMS and H&E) were then dehydrated using absolute ethanol, cleared in xylene and cover-slipped using xylene-based mounting medium. Unless specified otherwise, w/v and v/v solutions were prepared in dH_2_O and procedures were conducted at room temperature. Sample processing, sectioning and staining were performed by the Melbourne Histology Platform (The University of Melbourne).

Histology & qualitative slide analysis was performed for 5 larvae from each infection condition (6 hours 25°C PI, 6 hours 37°C PI, 48 hours 25°C PI or 48 hours 25°C PI) and for two larvae from each negative control condition (no injection 48 hours 25°C and no injection 48 hours 37°C) to ensure that observations were representative. All slides were examined using an Olympus BX40 microscope. High-power images were captured using an Olympus DP71 microscope camera and selected slides were digitised using a Pannoramic 480 digital scanner (3DHistech) for low-power images. Slide scanning was performed by the Phenomics Australia Histopathology & Slide Scanning Service (The University of Melbourne).

### Extraction and examination of larval haemolymph

Larvae that had not died within the designated incubation period (6 or 48 hours) were briefly washed in 70% ethanol to remove microorganisms adhered to the cuticle surface, dried with a clean paper towel, and chilled to 4°C. Haemolymph was then collected from individual larvae as follows: the dorsal side of each larva was bent over a P1000 micropipette tip taped onto a petri dish to expose the ventral surface and allow access to the sternites. The sternites were then gently pricked with a 25G needle to release a droplet of haemolymph. The haemolymph droplet (5-10 μl) was then carefully aspirated by micropipette and transferred to a microfuge tube. Haemolymph collected from 6 individual larvae was pooled and fixed for at least 30 minutes with 200 μl of 4% v/v paraformaldehyde (in PME buffer) at room temperature and centrifuged at 3000 g for 10 min. After centrifugation the supernatant was removed, and the haemocyte pellet was washed with 200 μl PME buffer. Samples were then centrifuged at 3000 g for 10 min, and the majority of the supernatant removed, leaving ~50 μl to resuspend the haemocytes. 5 μl of the haemocyte suspension was added to 5 μl CALC-DABCO mounting medium (20 ng/ml calcofluor white, 100 mg/ml DABCO in glycerol), gently mixed by pipette and mounted on a glass slide. Slides were examined using a ZEISS Axio Z1 microscope. Haemocytes were observed under DIC and calcofluor stained-fungal cells were observed using a 385 nm UV Colibri LED filter cube. Images were simultaneously captured in both channels on a ZEISS Axiocam 506 mono microscope camera. PME Buffer = 50 mM PIPES pH 6.7, 5 mM MgSO4, 25 mM EGTA pH 8.0.

## Data availability

The *T. marneffei* reference strain FRR2161 is publicly available from the American Type Culture Collection as ATCC18224 and the *niaD1ΔligD* laboratory strain (G821) will be distributed upon request. All data supporting the findings of this study are provided in the article itself or within the supplementary material.

## Acknowledgements

This research was supported by the Commonwealth through an Australian Government Research Training Program Scholarship [DOI: https://doi.org/10.82133/C42F-K220] awarded to B.E.W. The authors thank Minibeasts Enterprises for providing well-sized larvae for experiments, as well as the Melbourne Histology Platform (MHP) and the Phenomics Australia Histopathology & Slide Scanning Service at the University of Melbourne for use of their facilities and services. They are especially grateful to Laura Leone (MHP) for her efforts in optimising the larval fixation process.

## Notes

### Competing Interest Statement

The authors have declared no competing interest.

