## Supplementary Data for "A *Tenebrio molitor* model for *Talaromyces marneffei* infection and the importance of host cues for dimorphic switching"

### Supplementary material

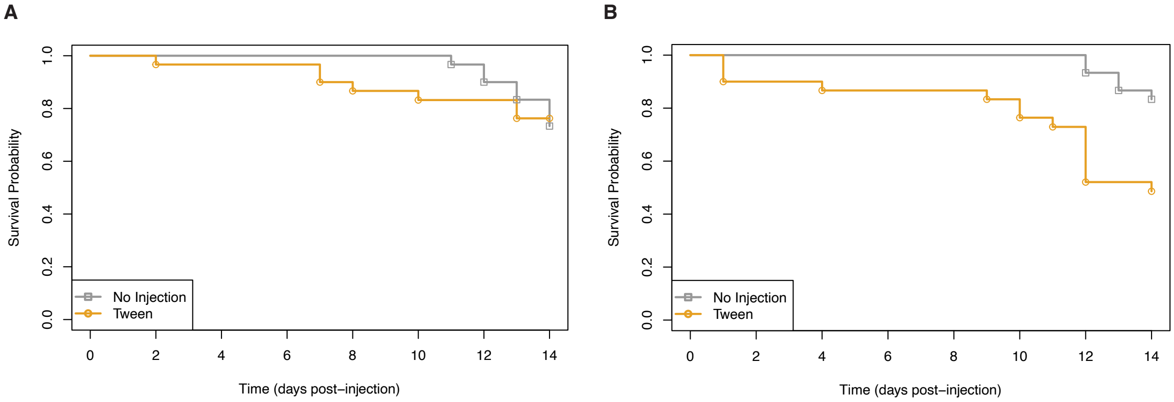


**Figure S1. Survival of *T. molitor* larvae at 37°C.** Kaplan-Meier plot showing survival of larvae that were not-injected (No Injection group) or injected with Tween solution and incubated at **(A**) 37°C or **(B)** 25°C. Experiments were performed with groups of 10 larvae and repeated three times. Survival was monitored for 14 days, and the results of the three replicates were then pooled together (n = 30) to build the survival curves. The early deaths (1-4 days PI) are presumed to be due to injection injuries at both temperatures.

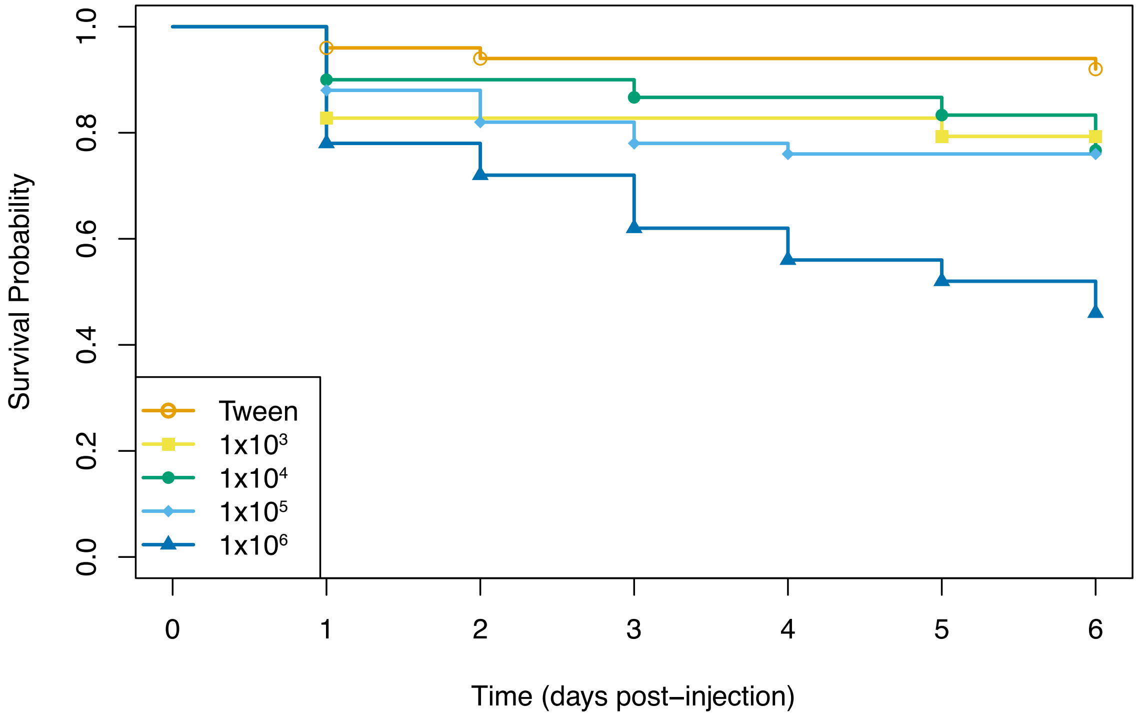


**Figure S2. Survival of *T. molitor* larvae infected with different concentrations of *T. marneffei* conidia at 37 °C.** Kaplan-Meier plots showing survival of larvae injected with Tween solution or different concentration of *T. marneffei* conidia. Survival of larvae injected with $10^3$ or $10^4$ conidia did not significantly differ from those injected with the Tween solution, while larvae injected with ≥ $10^5$ conidia did. Survival curves were built with data from 3-5 replicate experiments (3 for the $10^3$ or $10^4$ groups and 5 for all others) and pairwise LogRank test results are listed in Table 1.

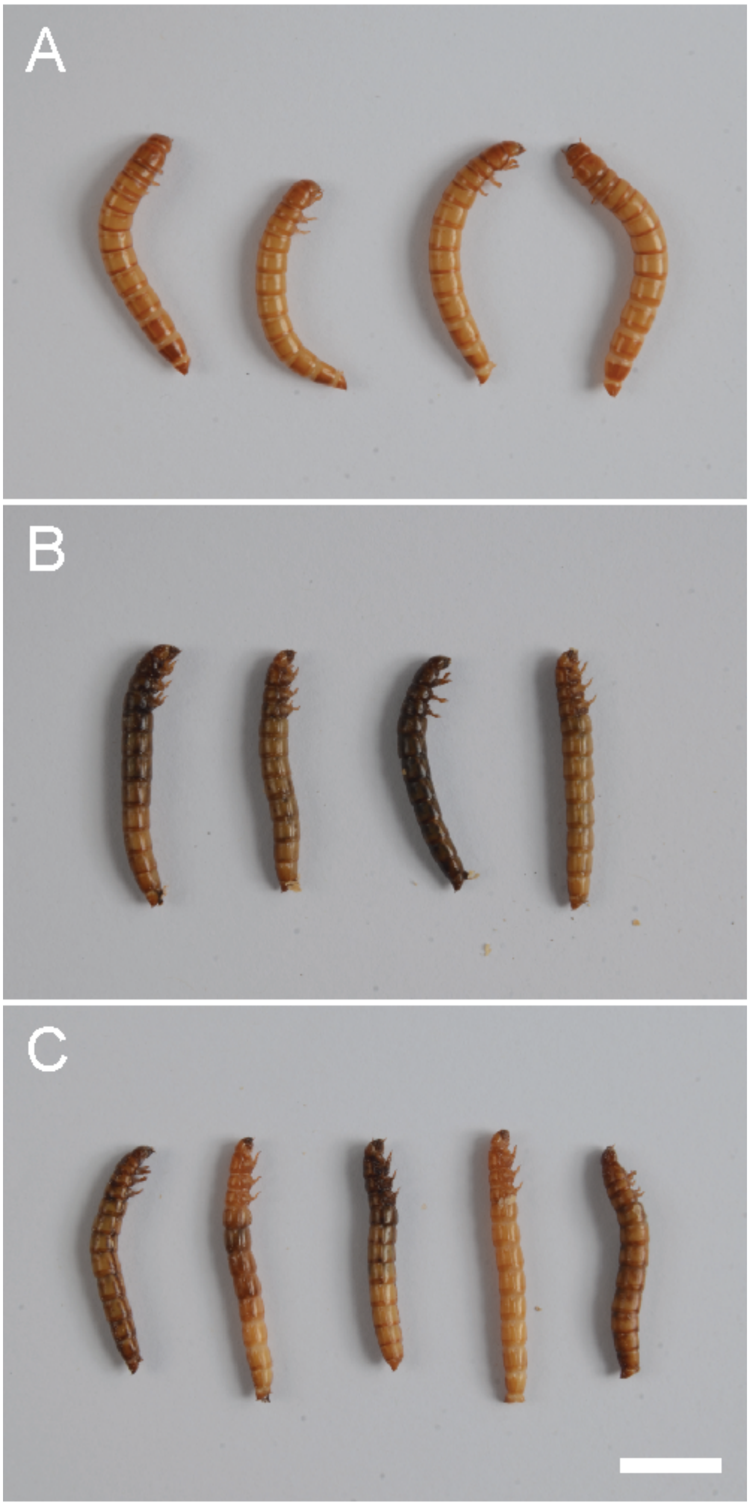


**Figure S3. Melanisation of *T. molitor* larvae.** Photographs of *T. molitor* larvae demonstrating differences in colour between healthy and dead larvae. **(A)** Live uninfected larvae (No Injection) without melanisation. **(B)** Dead larvae (4 days PI) that had been injected with $10^6$ conidia and incubated at 37°C **(C)** Dead larvae (1 day PI) that had been injected with $10^6$ conidia and incubated at 25°C. The amount of melanisation present when death was scored was highly variable, making it unsuitable as an indicator of mortality. Mortality was instead determined by response to physical stimuli. Scale bar = 1 cm in C (for A-C).

| Table S1. Presence of melanisation in infected and uninfected larvae incubated at 37 °C. | | | | |
|---|---|---|---|---|
| | **Dead (at time of death)** | | **Alive (at 6 days PI)** | |
| | **n** | **% melanised** | **n** | **% melanised** |
| **No Injection control** n= 20 | - | - | 20 | 5% |
| **Tween control** n= 20 | 2 | 100% | 18 | 38.9% |
| **Heat-Inactivated, $10^6$** n= 20 | 5 | 60% | 15 | 33.3% |
| **Viable, $10^5$** n= 20 | 3 | 83.3% | 17 | 29.4% |
| **Viable, $10^6$** n= 20 | 11 | 90.9% | 9 | 88.9% |

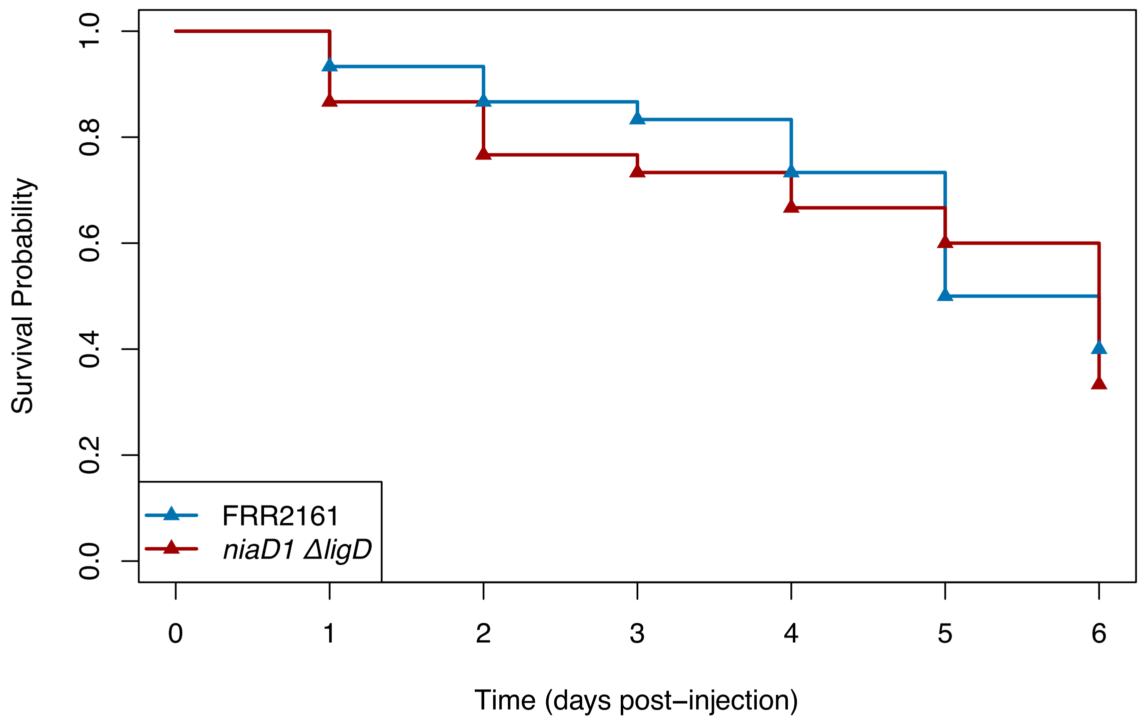


**Figure S4. Survival of *T. molitor* larvae infected with the NHEJ-deficient strain does not significantly differ from the reference strain.** Kaplan-Meier plots showing survival probabilities for larvae injected with $10^6$ *T. marneffei* conidia of the FRR2161 reference strain or the commonly used *niaD1DligD* laboratory strain over a 6-day period at 37°C. Pairwise LogRank test for differences in survival between the groups yielded a non-significant result (p-value = 0.7). Both survival curves were built with pooled data from 3 replicate experiments (n= 30 per group).

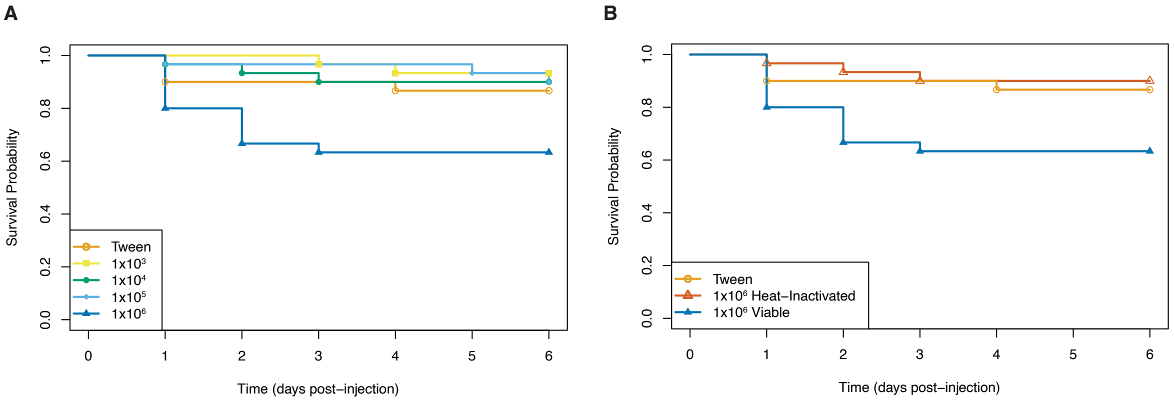


**Figure S5. Survival of *T. molitor* larvae infected with different concentrations of *T. marneffei* conidia at 25°C.** Kaplan-Meier plots showing survival probabilities for larvae injected with Tween solution or *T. marneffei* conidia over a 6-day period. **(A)** Injection with < $10^6$ conidia does not significantly impact survival compared to injection with Tween solution alone. Larvae injected with $10^6$ conidia did have significantly reduced survival compared to control groups. **(B)** Survival of larvae injected with $10^6$ Heat-Inactivated conidia does not significantly differ from those injected with the Tween solution, indicating the survival decline of larvae injected with $10^6$ live conidia is due to the presence of viable *T. marneffei*. All survival curves were built with pooled data from 3 replicate experiments (n= 30 per group).

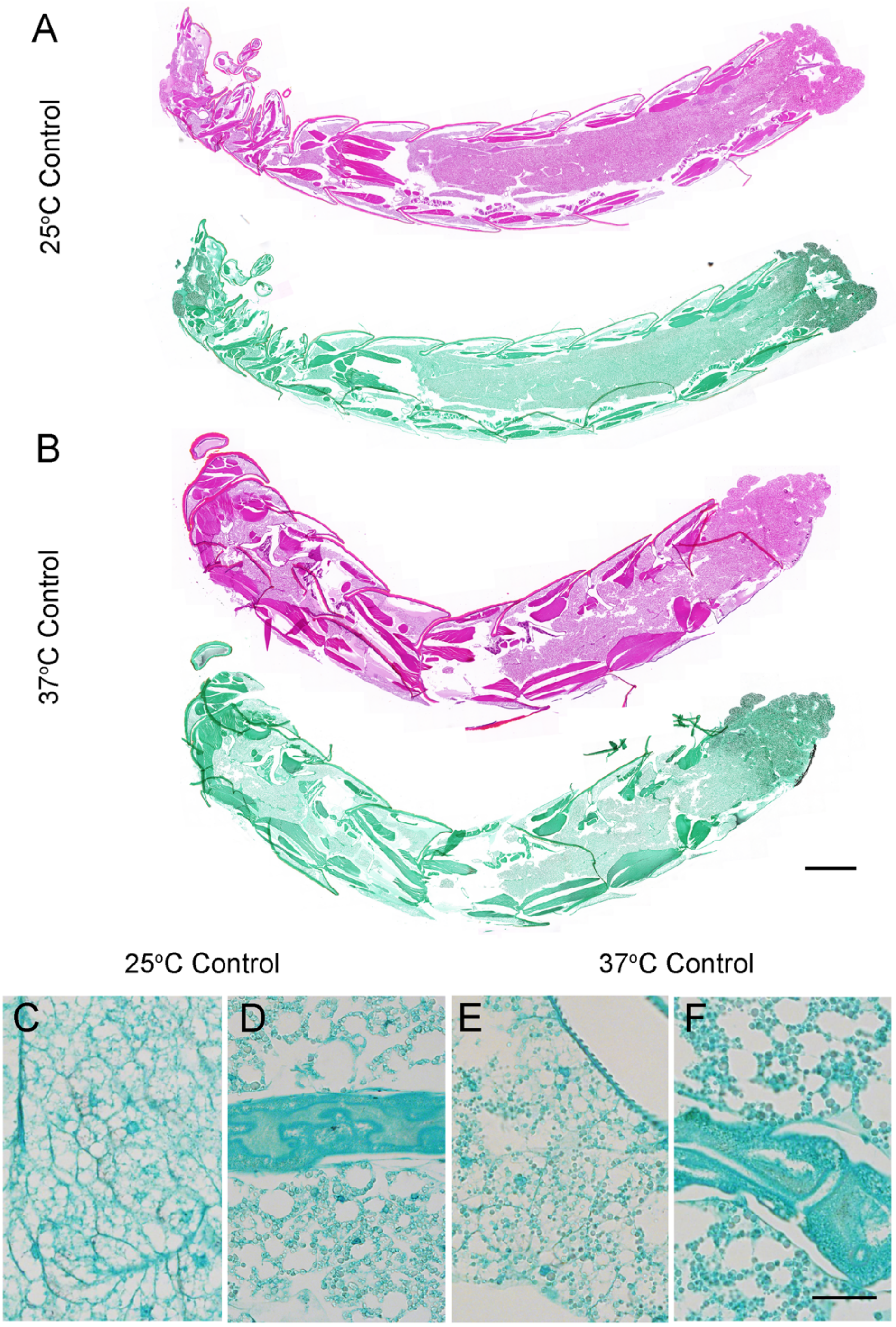


**Figure S6. Histology of NI control larvae.** (A, B) H&E-stained (pink) or GMS-stained sagittal sections of NI control larvae incubated at 25°C (A) or 37°C (B). Sagittal sections preserve spatial and tissue organisation revealing anterior (left) to posterior (right) morphology. (C – E) Higher power images of GMS-stained sections show examples of regions equivalent to those shown in Figure 3. No fungal staining was observed, which is consistent throughout all control sections. No other notable differences (e.g. tissue morphology) were observed between control and infected larvae (Figure 3). Scale bar = 1 mm in B (for A, B) and scale bar = 20 μm in E (for C – E).

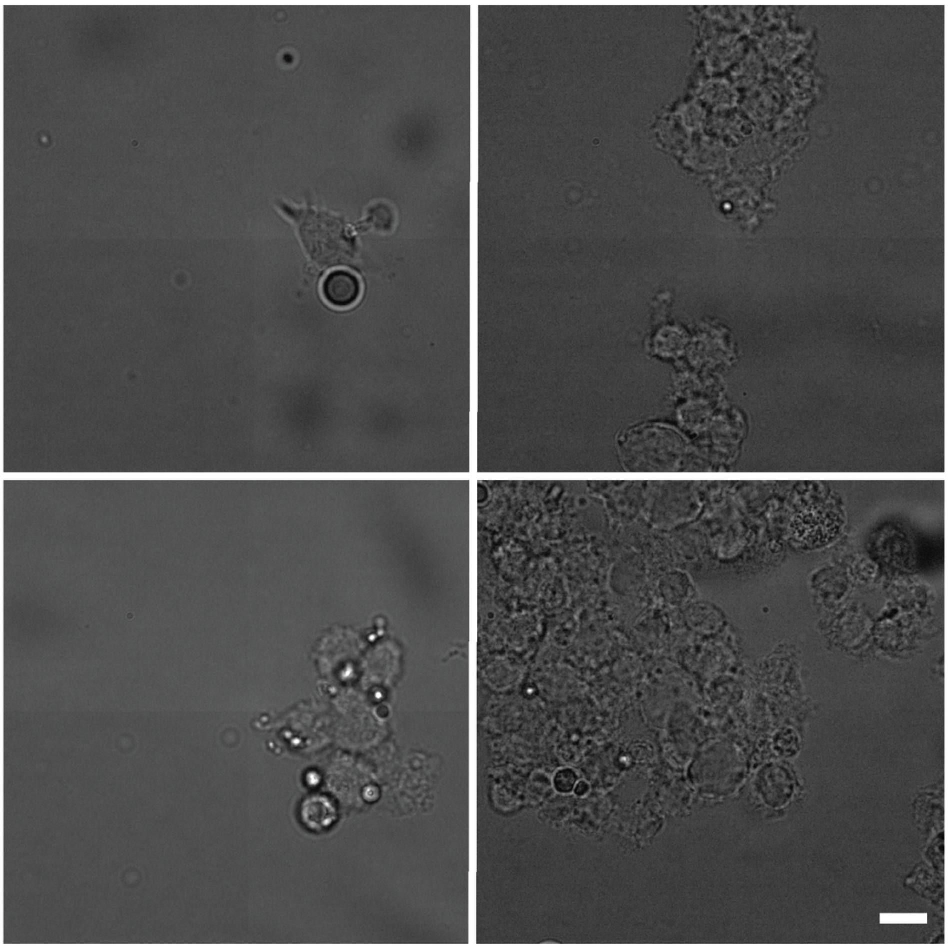

**Figure S7. Successful isolation of *T. molitor* haemocytes.** Representative micrographs of *T. molitor* haemocytes isolated from *T. molitor* larvae following fixation, washing and centrifugation steps (see Materials & Methods for details). Haemocyte suspensions encompassed single cells (top left), as well as small (bottom left, top right) and large (bottom right) cell aggregates. Images captured using brightfield microscopy. Scale bar = 10 µm and applies to all panels.

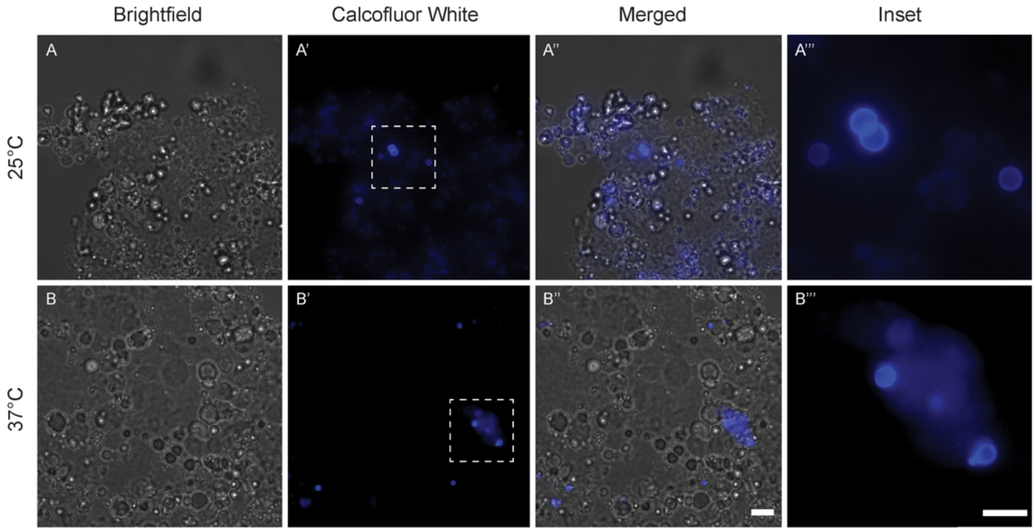


**Figure S8. Conidia present in haemolymph 6 hours post-injection.** Representative micrographs of *T. molitor* haemolymph collected from larvae incubated at 25 or 37°C for 6-hours post-injection with $10^6$ *T. marneffei* conidia and stained with 10 mg/ml calcofluor white to observe fungal cell walls. Conidia (Calcofluor White) are found in association with clusters of haemocytes (brightfield) at both temperatures. Scale bar in B'' (for A-A'', B'-B'') = 10 μm and Scale bar in B''' (for A''', B''') = 5 μm.

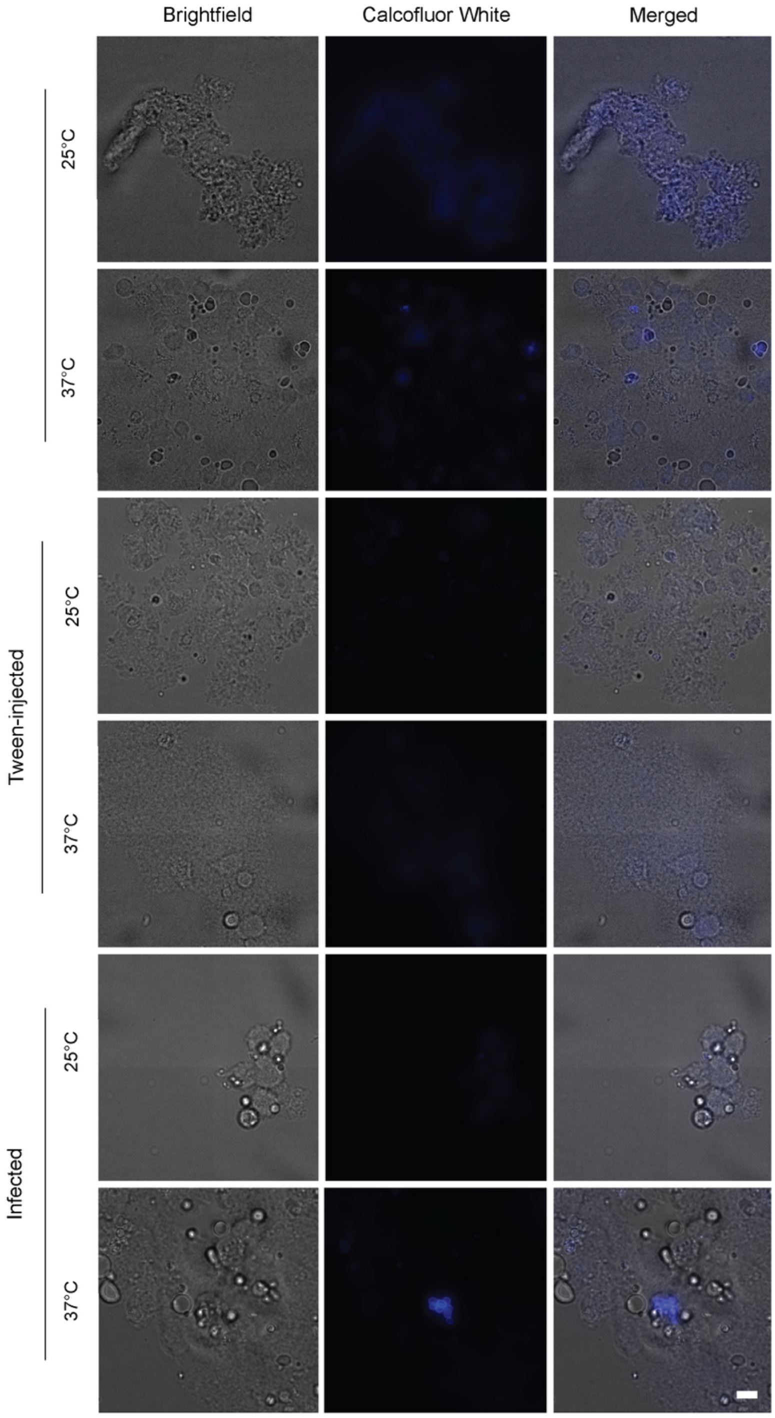


**Figure S9. Conidia present in haemolymph 6 hours post-injection**
Representative micrographs of *T. molitor* haemolymph collected from NI control, Tween control and *T. marneffei*-infected ($10^6$ conidia injected) following incubation at 25 or 37°C and stained with 10 mg/ml calcofluor white to observe fungal cell walls. No fungal cells are visible in NI control, Tween control, or 25°C-Infected samples, while distinct clusters of oval yeast cells (calcofluor white) in association with clusters of haemocytes were evident (visible under brightfield) in the or 37°C-Infected samples. Equivalent display settings were applied to all images to distinguish between background fluorescence and fungal cell staining. Scale bar = 10 μm and applies to all panels.

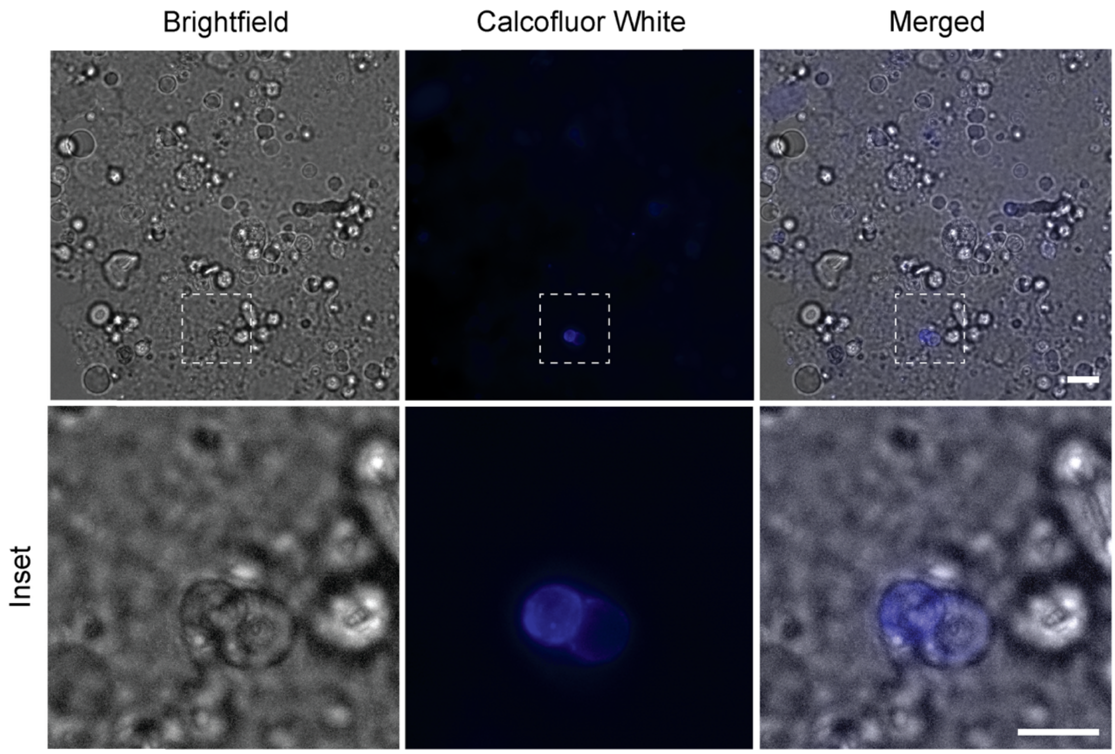


**Figure S10. Yeast-like cells in haemolymph 48 hours post-injection and incubation at 25°C.** Micrograph of *T. molitor* haemolymph collected from larvae incubated at 25°C for 48-hours post-injection with $10^6$ *T. marneffei* conidia and stained with 10 mg/ml calcofluor white to observe fungal cell walls. Two yeast cells (stained with calcofluor white and indicated with white arrowheads) are associated with a cluster of haemocytes (visible under brightfield). These were the only fungal cells identified in the 25°C-Infected-48-hour haemolymph samples. Scale bar for top row = 10 $\mu$m and scale bar for inset panels (bottom row) = 5 $\mu$m.

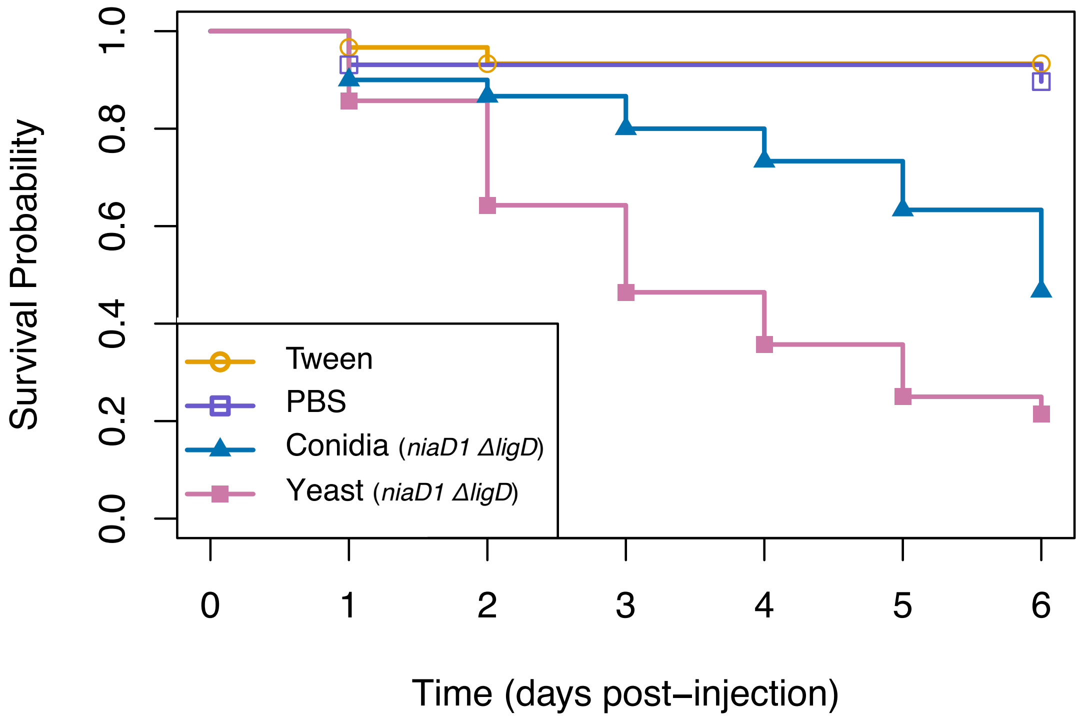


**Figure S11. Differences in survival of larvae infected with $10^6$ *niaD1DligD* conidia or *ex vivo* yeast cells at 37 °C.** Kaplan-Meier plot showing survival of larvae injected with viable *niaD1DligD* conidia in Tween, viable *ex vivo niaD1DligD* yeast cells in PBS, Tween solution and PBS (vehicle only controls), incubated at 37 °C. Direct injection with NHEJ-deficient yeast cells significantly increases the rate of survival decline compared to injection with NHEJ-deficient conidia (Table S2), as was observed for the FRR2161 reference strain (Fig. 5). There was no significant difference in survival of larvae injected with Tween solution or PBS, confirming that neither vehicle solution impacted survival outcomes. Survival curves were built with pooled data from three replicate experiments.

**Table S2. Pairwise LogRank test for differences in survival when injected with *niaD1 ΔligD* conidia or yeast**

| | **Viable Conidia, $10^6$** n= 30 | **PBS control** n= 29 | **Tween control** n= 30 |
|---|---|---|---|
| **PBS control** n= 29 | N/A | - | 0.6 |
| **Viable Conidia, $10^6$** n= 30 | - | N/A | 7e-05 |
| **Viable Yeast cells, $10^6$** n= 28 | 0.02 | 6e-07 | N/A |

Dashes (-) denote self-comparisons and N/A denotes group combinations which are not directly comparable.

| Table S3. Viable counts of conidia & yeast cell suspensions when plating for ~ 100 colonies | | | | |
|---|---|---|---|---|
| | **Suspension** | **Mean (Viable count)** | **Standard Deviation** | **Yeast : Hyphae ratio** |
| **FRR2161** | Conidia | 44 | ±15.59 | N/A |
| | Yeast | 19 | ±0.58 | 2:1 |
| ***niaD1 ΔligD*** | Conidia | 30 | ±9.82 | N/A |
| | Yeast | 16 | ±6.35 | 2:1 |
